# Developmental chromatin programs determine oncogenic competence in melanoma

**DOI:** 10.1101/2020.05.09.081554

**Authors:** Arianna Baggiolini, Scott J. Callahan, Tuan Trieu, Mohita M. Tagore, Emily Montal, Joshua M. Weiss, Sam E. Tischfield, Yujie Fan, Nathaniel R. Campbell, Nathalie Saurat, Travis Hollmann, Theresa Simon-Vermot, Satish K. Tickoo, Barry S. Taylor, Richard Koche, Ekta Khurana, Lorenz Studer, Richard M. White

## Abstract

Oncogenes are only transforming in certain cellular contexts, a phenomenon called oncogenic competence. The mechanisms regulating this competence remain poorly understood. Here, using a combination of a novel human pluripotent stem cell (hPSC)-based cancer model along with zebrafish transgenesis, we demonstrate that the transforming ability of BRAF^V600E^ depends upon the intrinsic transcriptional program present in the cell of origin. Remarkably, in both systems, melanocytes (MC) are largely resistant to BRAF. In contrast, both neural crest (NC) and melanoblast (MB) populations are readily transformed. Molecular profiling reveals that NC/MB cells have markedly higher expression of chromatin modifying enzymes, and we discovered that the chromatin remodeler ATAD2 is required for response to BRAF and tumor initiation. ATAD2 forms a complex with SOX10, allowing for expression of downstream oncogenic programs. These data suggest that oncogenic competence is mediated by developmental regulation of chromatin factors, which then allow for proper response to those oncogenes.

## Main Text

Alterations to DNA structure, including mutations, copy number variations and gene fusions are typically considered initiating events in most human cancers (*1*). However, these alterations are layered onto existing transcriptional programs in the cell of origin. The importance of these pre-existing cellular lineage programs is highlighted by the fact that certain DNA mutations are only tumorigenic in certain cell types (*2, 3*), which we refer to as oncogenic competence.

In melanoma, it has been shown that tumor initiation by BRAF^V600E^ activates a neural crest lineage program (*4, 5, 6, 7*). In particular *crestin*, a gene that is specifically expressed in neural crest cells and is downregulated later during embryonic development, is reactivated in melanoma initiating cells and then maintained in the tumor (*6*). The activation of NC-lineage specific mechanisms (*4-10*) together with oncogene mutations such as BRAF^V600E^ are fundamental for the acquisition of a malignant state (*11, 12*). However, it is not known why a NC-like state is required and particularly susceptible to oncogenic transformation by BRAF and what the factors are regulating this state.

Lineage programs are centrally intertwined with the cell of origin. Within a developmental lineage, cells can exist along a wide spectrum of differentiation states. After emergence from the neural crest, migrating neural crest progenitors give rise to populations of melanoblasts, melanocyte stem cells, or differentiated melanocytes. Which cell along that spectrum is intrinsically capable of giving rise to melanoma even above and beyond the influence of extrinsic microenvironmental factors has been a matter of debate (*13, 14, 15*). Here, we asked what mechanisms define melanoma competence and how it is regulated.

### Zebrafish models show that NC and MB, but not MC, are oncogenic competent

The melanocyte lineage develops as a hierarchy of cells that start as undifferentiated neural crest cells, proceeding through a melanoblast stage and then finally differentiating into mature, pigmented melanocytes. To understand which cells within this lineage are most sensitive to an oncogenic insult, we engineered zebrafish to initiate tumors in either neural crest (NC), melanoblasts (MB) or melanocytes (MC) by using stage-specific promoters to drive BRAF^V600E^. We used the *sox10* promoter to drive BRAF^V600E^ expression in the NC, the *mitfa* promoter to drive it in the MB and the *tyrp1* promoter to drive it in the MC. These transgenic constructs were injected into 1-cell embryos of a p53^-/-^ deficient background, and the animals were allowed to grow up to adulthood to monitor for tumor formation (Fig. 1A). We found that animals that expressed the BRAF^V600E^ oncogene either in the NC cells or the MB developed aggressive tumors (Fig. 1B, 1C, 1D, fig. S1A and S1B) with nearly 100% penetrance in the NC lineage (Fig. 1B). Surprisingly, the *tyrp1*:BRAF^V600E^;p53^-/-^ transgenic animals failed to develop tumors, but instead developed small patches of nevus-like cells (Fig. 1B, 1E, 1H, 1K and fig. S1C). We analyzed both the NC- and the MB-derived tumors and found that they stained equally for pERK (Fig. 1I, 1J), indicating that the BRAF pathway (Fig. 1F, 1G) was being activated. H&E showed that the NC- and MB-derived tumors were histologically distinct (fig. S1A, S1B). The NC-derived tumors appeared undifferentiated and with little melanin. Immunohistochemistry showed that the NC-derived tumors were predominantly positive for the neuronal markers huc/hud and ncam and sparsely for sox10 (Fig. 1L, 1N, 1P), likely reflecting the multipotency of the NC (*16, 17, 18*) and resembling reports of melanoma with neuronal type differentiation (*19, 20*). On the contrary, the MB-derived tumors had an appearance characteristic of typical cutaneous melanoma, with numerous pigmented areas, and they stained positive for not only sox10 (Fig. 1M), as previously shown for most melanomas (*8, 21*), but also positive for mlana (Fig. 1Q), while being negative for the neuronal markers huc/hud (Fig. 1O). To further confirm the differences between the NC- and MB-derived tumors, we performed RNA-seq and found that these tumors clustered separately from each other by PCA and hierarchical clustering (Fig. 1R, fig. S1D and table S1). Examination of specific pathways in each tumor revealed that, consistent with our immunohistochemistry, the NC-derived tumors expressed neuronal genes while the MB-derived melanomas expressed genes related to the melanocytic lineage (fig. S1E). We performed Gene Set Association Analysis (GSAA) and confirmed that NC-derived tumors displayed a strong neuronal phenotype and a gene signature that characterizes poor survival in neuroblastoma (fig. S1F) (*22*). These data suggest that competence to respond to BRAF^V600E^ is biased towards cells of origin exhibiting either a NC or MB gene program, that these give rise to distinct tumors, and that MC are relatively resistant to this insult.

**Fig. 1.**
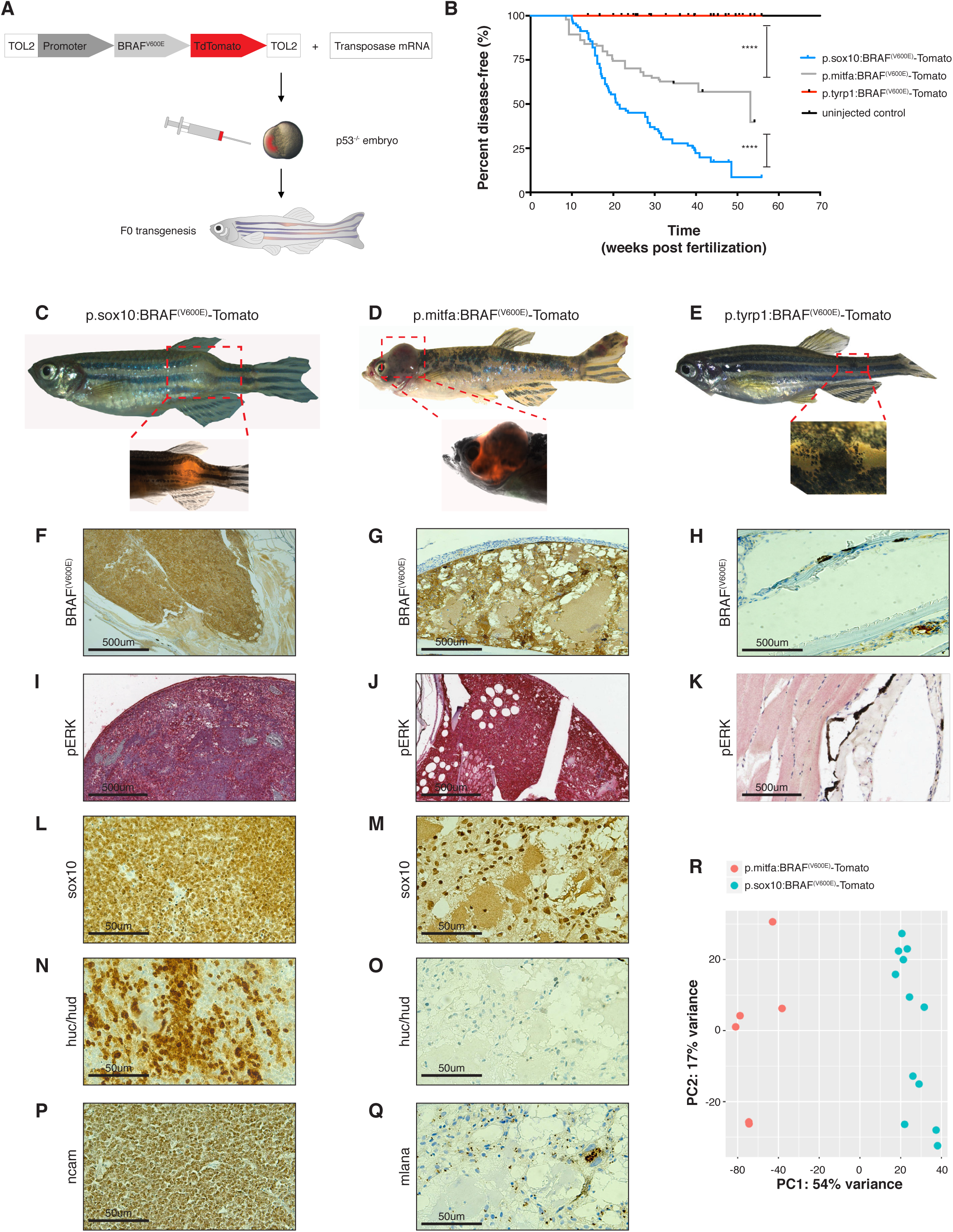
Zebrafish models show that the NC and MB states, but not the MC state, are cancer competent. **(A)** Schematic drawing of zebrafish F0 transgenesis. F0 zebrafish transgenic fish were engineered by injection of p53^−/−^ single-cell embryos with transposase mRNA together with TOL2 flanked plasmids, which encoded a stage-specific promoter (*sox10, mitf, tyrp1*) driving BRAF^(V600E)^ fused to TdTomato. **(B)** Kaplan-Meier curves of F0 p53^-/-^ transgenic zebrafish injected with plasmids driving BRAF^(V600E)^ fused to TdTomato under either the NC-specific promoter *sox10* (n=92 biological replicates), the MB-specific promoter *mitfa* (n=94 biological replicates), or the MC-specific promoter *tyrp1* (n=49 biological replicates) or uninfected control (n=86 biological replicates). **** = p < 0.0001 for the comparison of the tumor-free survival curves of fish with MB-derived tumors and MC-derived nevus-like structures; **** = p < 0.0001 for the comparison of the tumor-free survival curves of fish with NC- and MB-derived tumors; log-rank (Mantel-Cox) test. **(C)** Representative NC-derived tumor developed in the *sox10*:BRAF^(V600E)^ p53^−/−^ transgenic fish. **(D)** Representative MB-derived tumor developed in the *mitfa*:BRAF^(V600E)^ p53^−/−^ transgenic fish. **(E)** Representative nevus-like structure developed in the *tyrp1*:BRAF^(V600E)^ p53^−/−^ transgenic fish. (**F−K)** Immunohistochemistry for BRAF^(V600E)^ and phosphoERK in the NC- and MB-derived tumors and in the MC-derived nevus-like structure. (**L−Q)** Immunohistochemistry staining for sox10, huc/hud, ncam and mlana. NC-derived tumors were positive for the neuronal marks huc/hud and ncam **(N, P)** and mostly negative for sox10 expression **(L)**. MB-derived tumors were melanomas positive for sox10 **(M)**, mlana **(Q)**, and negative for the neuronal marks huc/hud **(O)**. **(R)** PCA plot of *mitfa*-derived tumors (n=6, M, red) and *sox10*-derived tumors (n=12, S, blue) for whole genome RNA-seq demonstrated a clear separation at the transcriptional level.

### A hPSC-based cancer model recapitulates the zebrafish models

Because mouse studies routinely use MC promoters to efficiently drive melanoma (i.e. the tyr-Cre line, *23*), this raised the question of whether the above findings were unique to the zebrafish system. To interrogate this, we developed a novel human cancer paradigm based on the use of hPSCs to model oncogenic competence for melanoma in a manner similar to the zebrafish studies. We previously demonstrated that hPSCs can be progressively differentiated into NC cells, MB or MC (*24*). We used gene targeting in hPSCs to introduce oncogenic BRAF^V600E^ and to inactivate the tumor suppressors RB1, TP53 and P16 (referred to hereafter as 3xKO cells) (Fig. 2A). These 3xKO engineered cells were then differentiated into NC cells, MB and mature MC (*24, 25*) (Fig. 2A and fig. S2A, S2B) and BRAF^V600E^ induced by doxycycline (fig. S2L). Western blot analysis for pERK demonstrated equal activation of the BRAF pathway across all three cell types (Fig. 2B), similar to the results in the zebrafish studies above. To test their tumorigenic capacity, we transplanted each of three cell types subcutaneously in immunodeficient NOD-*scid* IL2Rgamma^null^ (NSG) mice, a model that has been used before to assess the tumorigenic potential of the cells (*26, 27*), and the mice were exposed either to a dox-containing or to a normal diet. Similar to the zebrafish, we found that both NC cells and MB expressing BRAF^V600E^ in 3xKO background readily formed tumors in the mice (Fig. 2C, 2D), but that the 3xKO MC largely failed to do so, with only a single animal developing a tumor under this condition (Fig. 2E). Residual cells from the site of MC transplant at day 197 post injection showed features of a benign nevus-like structure consistent with them becoming senescent over time (fig. S2K). As a control, we also transplanted wildtype (WT) NC cells, MB and MC and found that these were unable to grow in vivo, as expected (data not shown). We performed histological analysis of the NC- and MB-derived tumors (Fig. 2F-S and fig. S2C-J) and found high level expression of BRAF^V600E^ and pERK in both (Fig. 2F-2I). Analogous to the zebrafish tumors, the human PSC-based 3xKO NC cells gave rise to tumors that showed a strong preponderance of neuronal markers (Fig. 2P, 2R), whereas the 3xKO MB were positive for all the common markers of melanoma and were pathologically categorized as desmoplastic melanomas (Fig. 2K, 2M, 2O). We performed Gene Set Enrichment Analysis (GSEA), and we found that NC cells were transcriptionally similar to the NC-derived tumors in the fish (Figure S3B) and that MB were transcriptionally similar to the MB-derived tumors in the fish (fig. S3C), which further corroborated the comparability between the zebrafish and the hPSC-derived cancer models. To ensure that our hPSC-derived tumors are relevant to human patients, we performed RNA-seq of the BRAF^V600E^ 3xKO NC cells, MB and MC and compared their expression profiles to data from The Cancer Genome Atlas (TCGA), using a published signature for melanoma subgroups (*28*). Strikingly, we observed that the hPSC-derived 3xKO dox NC cells and 3xKO dox MB strongly clustered with the human melanoma patient samples (Fig. 2T), whereas the hPSC-derived 3xKO dox MC did not. Interestingly, we unexpectedly found that the 3xKO NC cells and 3xKO MB, even without BRAF^V600E^ induction, could form tumors in mice indicating that loss of tumor suppressors alone gives these cells enough of a proliferative advantage to grow in this context. As mentioned, we did find a single animal that grew a tumor after transplant with 3xKO MC, but exome sequencing showed that this single tumor had gained copy number alterations in MYC (padj=2.30E^-15^) (data not shown), among other genomic changes, likely contributing to its capacity to grow in vivo. Thus, similar to what we found in the zebrafish, tumorigenic capacity in our novel human melanoma model is strongly biased towards NC or MB lineage, and that MC are relatively refractory to tumorigenesis although can be coaxed to do so with additional genetic lesions.

**Fig. 2.**
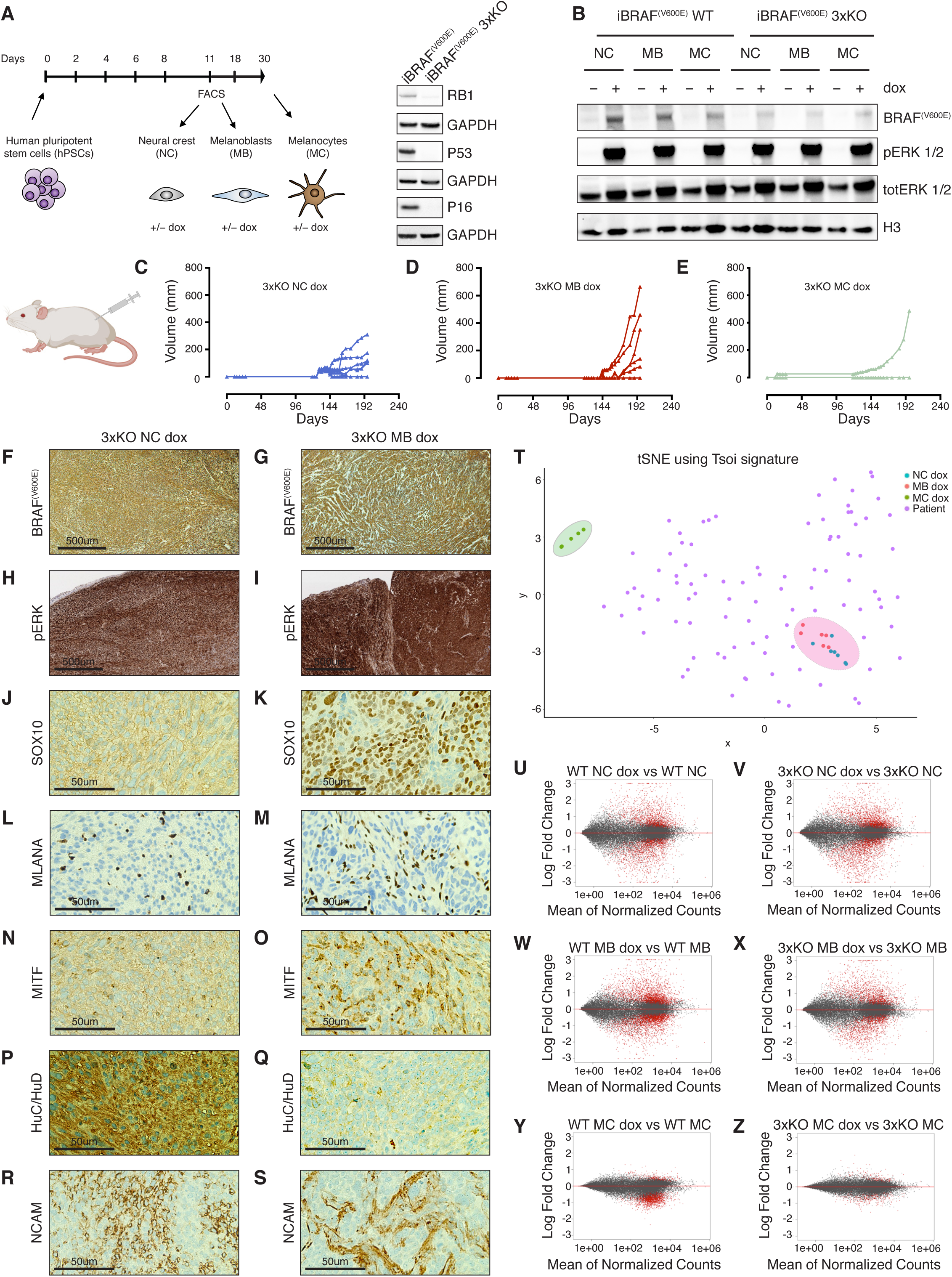
A hPSC-based cancer model recapitulates the zebrafish models and demonstrates that human NC and MB states are cancer competent, while the differentiated MC state is not. **(A)** Schematic summary of hPSCs differentiation into NC cells, MB and MC and Western blot of the dox-inducible BRAF^(V600E)^ (iBRAF^(V600E)^) hPSC line knockout for RB1, P53 and P16 (3xKO) using CRISPR/Cas9 technology. **(B)** Western blot of NC cells, MB and MC differentiated from either the iBRAF^(V600E)^ WT or the iBRAF^(V600E)^ 3xKO hPSCs. The cells were exposed to dox (1μg/ml) for 72h. (**C-E)** In vivo growth curves of 3xKO NC cells + dox (n=6 per group) **(C)**; in vivo growth curves of 3xKO MB + dox (n=6 per group) **(D)**. 3xKO MC + dox were not able to grow in vivo (n=6 per group, 1 outlier) (**E)**, but gave rise to nevus-like structures (Figure S2K). hPSCs-derived cells were injected subcutaneously in immunodeficient NSG mice exposed to a dox-containing diet. (**F**-**S)** Immunohistochemistry of NC-derived and MB-derived tumors + dox treatment. NC-derived tumors were undifferentiated and heterogeneous tumors, with strong neuronal features **(P, R)**. MB-derived tumors were diagnosed as melanomas and they were positive for all the common melanocytic marks **(K, M, O)**. (**T)** t-distributed Stochastic Neighbor Embedding (t-SNE) of 3xKO + dox NC cells, MB and MC samples and the TCGA melanoma samples using the Tsoi signature for melanoma subtypes. (**U**-**Z)** MA plots of the RNA-seq of WT NC cells, MB and MC ± dox treatment and 3xKO NC cells, MB and MC ± dox treatment (n=3 per condition). The mean of normalized counts of each gene was plotted against the log fold change following dox-induced BRAF^(V600E)^ expression within that condition. Adjusted p value cut-off of 0.05 was used for significantly differentially expressed genes (red).

### NC/MB cells have strong transcriptional responses to BRAF, but MC have little, despite comparable activation of pERK

To gain insight into why these cells differed in oncogenic competence, we performed RNA-seq of NC, MB and MC cells ± BRAF^V600E^ on both the WT and 3xKO background. We observed that dox-induced BRAF^V600E^ expression caused dramatic transcriptional changes in both the NC and MB (Fig. 2U, 2V, 2W, 2X, fig. S3A and S3D). In contrast, the transcriptional response to BRAF in MC was nearly absent, with only few genes being altered (23 genes upregulated (padj < 0.01) and 114 downregulated (padj < 0.01)) (Fig. 2Y, 2Z, fig. S3A, S3D and table S2). Thus, despite equally robust activation of pERK across all three cell types (as above in Fig. 2B), this indicates that the MC state was refractory to eliciting a transcriptional response following oncogene activation. This refractoriness to BRAF was not because the MC were post-mitotic, as proliferation of the 3xKO ± dox MC was comparable to the proliferation of 3xKO NC cells (see Fig. 5G). Because pERK ultimately acts to promote tumorigenesis via activation of downstream transcriptional responses, it appears that the failure of MC to be transformed is likely related to a failure of BRAF-dependent transcriptional induction in this lineage. This raised the question of what was intrinsically different between these cell types. To address this, we performed RNA-seq on WT NC, WT MB and WT MC (i.e. no transgenes) and analyzed the differentially expressed genes by GSEA pathway analysis (table S2). Amongst the most enriched pathways that separated the NC/MB cells from the MC was the GO term Chromatin modification (NES= 1.901, FDR= 0.06, Fig. 3A). In examining individual genes which were enriched in these stages, we found a striking enrichment for many chromatin modifier genes, including readers (e.g. ATAD2, CHD9, BAZ1A), writers (e.g. EZH2, PRMT8, HDAC9) and erasers (e.g. JMJD1C, TET1, TET2), all of which were markedly higher in the NC/MB cells compared to the MC (Fig. 3B and 3C). This suggested the possibility that the NC/MB cells might have intrinsically higher capability to rewire their chromatin state in response to BRAF^V600E^ and render them competent for melanoma initiation.

**Fig. 3.**
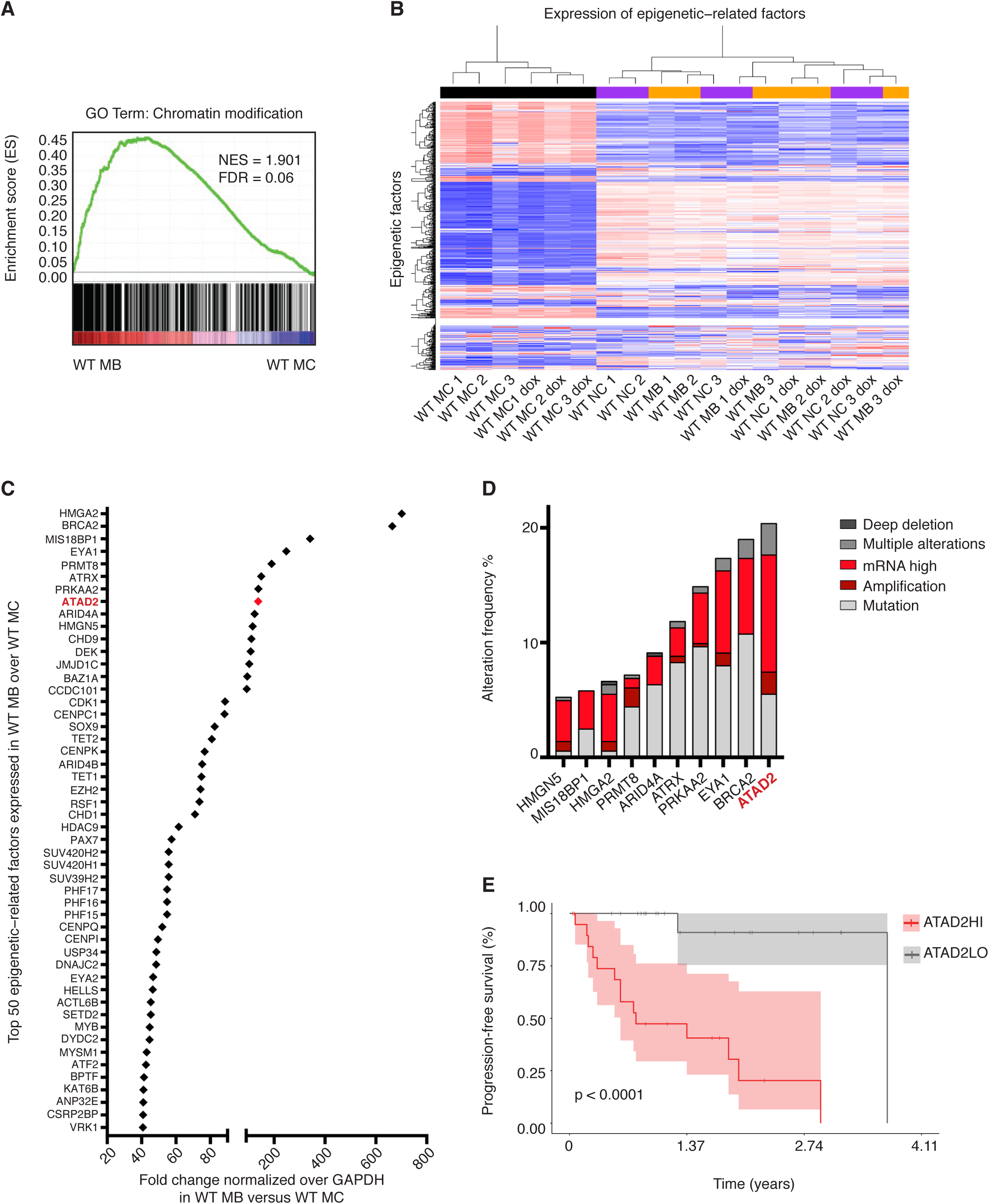
Cancer competence is reflected by a distinct chromatin landscape and ATAD2 is a key chromatin modifier shared between human NC/MB cells and patient melanoma cells. **(A)** GSEA comparing WT MB to WT MC identified chromatin modification as the pathway most enriched in WT MB. **(B)** Unsupervised clustering of WT ± dox NC cells (purple), MB (orange) and MC (black) depending on the expression profile of epigenetic-related factors. WT ± dox MC showed a distinct profile from the one of the WT ± dox NC cells and MB. **(C)** Top 50 epigenetic-related factors expressed in WT MB and downregulated in the differentiated WT MC. **(D)** Alteration frequency of the top 10 epigenetic-related factors in TCGA SKCM melanoma patient samples. **(E)** Kaplan-Meier overall survival curve of TCGA SKCM patients belonging either to the ATAD2HI or to the ATAD2LO group with log-rank p value reported.

**Fig. 4.**
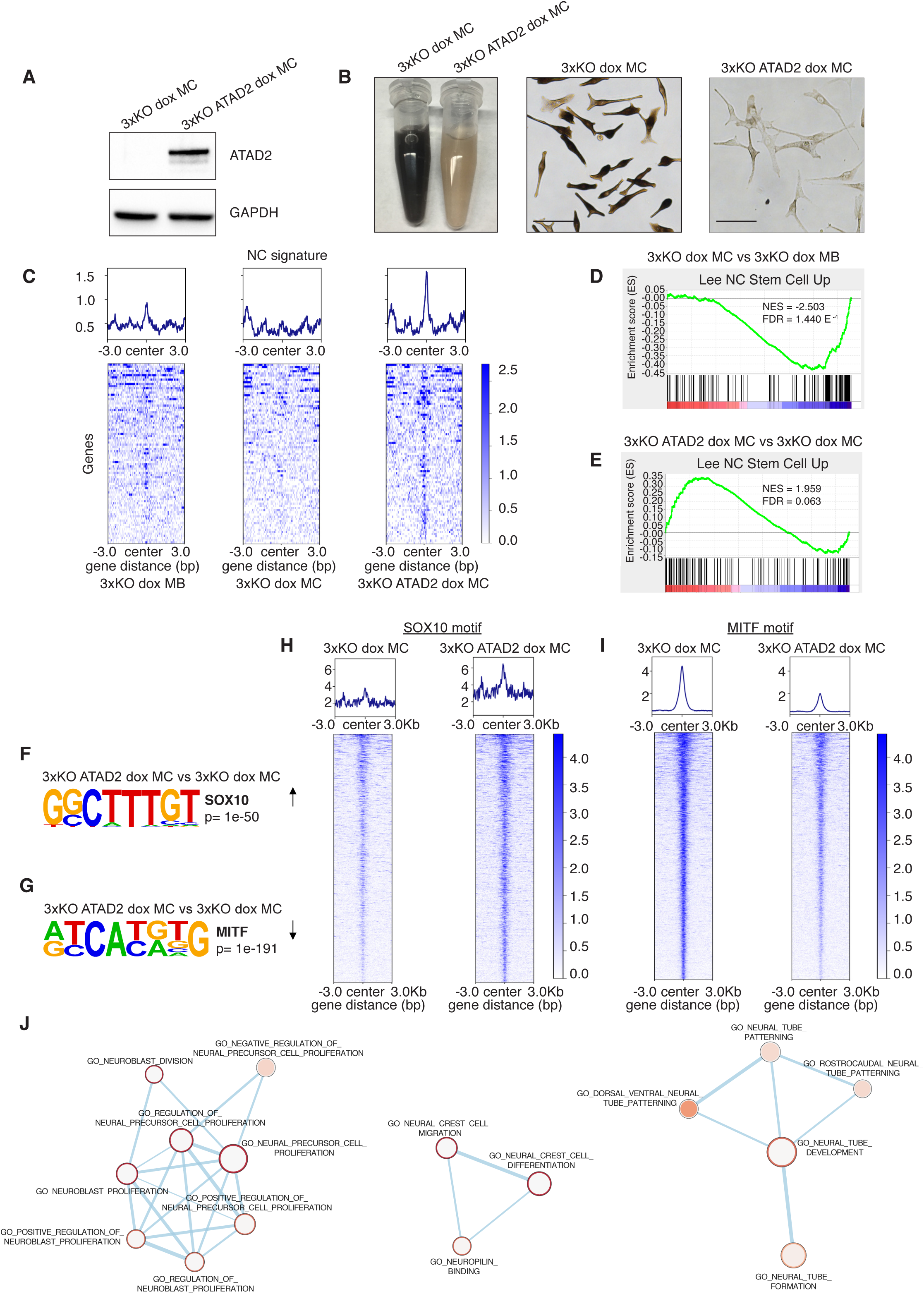
ATAD2 expression in MC reshapes the chromatin around NC/MB loci and reactivates a developmental signature. **(A)** Western blot for lenti-induced ATAD2 expression in 3xKO dox MC. **(B)** Representative pictures of 3xKO dox MC, on the right side, and of 3xKO ATAD2 dox MC, on the left side. Scale bars: 50 μm. **(C)** Tornado plots of the GSEA of the ATAC-seq for genes belonging to the NC signature in 3xKO dox MB, 3xKO dox MC and 3xKO ATAD2 dox MC. **(D-E)** GSEA of the ATAC-seq of 3xKO dox MC compared to 3xKO dox MB **(D)** and of 3xKO ATAD2 dox MC compared to 3xKO dox MC **(E)**. **(F)** Homer motif discovery shows that the SOX10 motif is one of the most enriched motifs (p value < 1e^−50^) in 3xKO ATAD2 dox MC compared to 3xKO dox MC. **(G)** Homer motif discovery shows that the MITF motif is the most closed motif (p value < 1e^−191^) in 3xKO ATAD2 dox MC compared to 3xKO dox MC. **(H)** Tornado plots depict the opening of the chromatin specific for the SOX10 motif. **(I)** Tornado plots depict the closure of the chromatin specific for the MITF motif. **(J)** Network analysis of the genes with increased accessibility for the SOX10 binding motif in 3xKO ATAD2 dox MC compared to 3xKO dox MC.

**Fig. 5.**
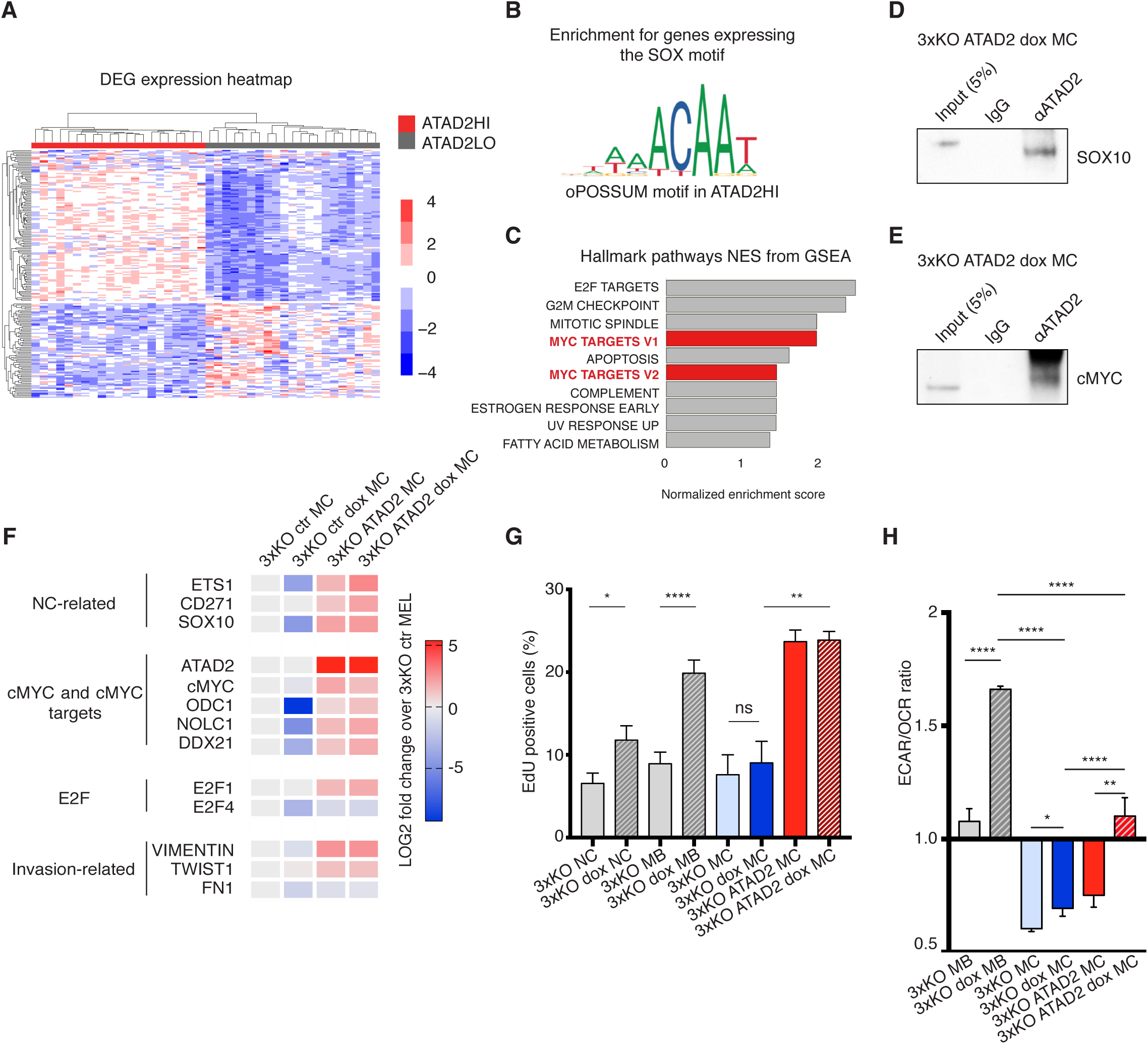
ATAD2 promotes melanoma phenotypes through cMYC and SOX10 in both clinical samples of cutaneous melanoma and in the hPSC-derived cancer model. **(A)** Heatmap plot of differentially expressed genes (DEG) in the ATAD2HI patient group versus the ATAD2LO patient group. **(B)** Identification of the SOX binding motif on genes co-expressed in the ATAD2HI patient groups, determined by analysis with the oPOSSUM software tool. **(C)** Top 10 hallmark pathways from GSEA enriched in the ATAD2HI patient group compared to the ATAD2LO patient group. **(D-E)** Co-IP analysis of protein lysates of 3xKO ATAD2 dox MC using either the ATAD2 or the control IgG antibody and then blotted against SOX10 **(D)** and cMYC **(E)**. **(F)** qRT-PCR of a subset of NC-related, cMYC-related and invasion-related genes in 3xKO dox MC, 3xKO ATAD2 MC and 3xKO ATAD2 dox MC. Heatmap depicts gene expression changes. n = 3 biological replicates. Significance illustrated in the figure refers to comparison to the control 3xKO MC. **(G)** EdU FACS analysis of 3xKO ± dox NC cells, 3xKO ± dox MB, 3xKO ± dox MC and 3xKO ATAD2 ± dox MC upon 30 min of EdU pulse. Data are shown as mean ± SEM, n=3. * = p < 0.05; ** = p < 0.005; **** = p < 0.0001. **(H)** Ratio between the OCR and the ECAR values of 3xKO ± dox MB, 3xKO ± dox MC and 3xKO ATAD2 ± dox MC. Data are shown as mean ± SEM, n=3. * = p < 0.05; ** = p < 0.005; **** = p < 0.0001.

### ATAD2 is a key chromatin modifier shared between NC/MB cells and melanoma cells

To identify which of these chromatin factors is likely most important in establishing competence, we analyzed the top 50 epigenetic-related factors that are higher in MB compared to MC (Fig. 3C) and then asked which of these is most commonly amplified or overexpressed in the human melanoma TCGA cohort. This analysis demonstrated that the top gene was ATAD2, an ATPase- and bromodomain-containing protein (*29*), which was altered in 20.4% of melanoma patients (Fig. 3D). We further asked if expression of ATAD2 correlated with patient survival and found that patients in the highest 20% of expression had significantly worse survival compared to the remaining patients (Fig. 3E and fig. S6A). Although there is no available information about ATAD2 in neural crest or melanoma development, these data nominated it as a key factor for promoting oncogenic competence in the NC/MB lineage.

### ATAD2 acts to reshape chromatin around key NC/MB loci

We wished to determine if ATAD2 was required for establishment of the NC/MB state and subsequent tumorigenesis. We generated a lentivirus that induced ATAD2 expression in the 3xKO MC (Fig. 4A) to a level comparable to what would be found in the NC cells or MB (fig. S4A). While MC without ATAD2 are deeply pigmented with melanin, reflecting their differentiated state, we noted that the 3xKO MC expressing ATAD2 lost their pigmentation (Fig. 4B). This suggested that ATAD2 expression leads to a dedifferentiated state possibly via affecting chromatin accessibility of NC-related genes in MC (*30-33*) (Fig. 4B). To test this idea, we performed ATAC-seq on 3xKO dox MB, 3xKO dox MC and 3xKO ATAD2 dox MC to assess global changes in chromatin accessibility. While MC had generally more open chromatin compared to MB, the addition of ATAD2 to the MC did not lead to a global increase in open chromatin (fig. S4B and S4C). Instead we found that overexpression of ATAD2 in the MC led to a significant increase in chromatin accessibility specifically at NC-related loci, to a level comparable if not greater than that found in the 3xKO dox MB themselves (Fig. 4C). GSEA analysis of the affected loci in the 3xKO ATAD2 dox MC versus the 3xKO dox MC showed a significant enrichment of NC-related genes (Fig. 4D, 4E). To gain insight into the transcription factors that are binding to these newly opened chromatin regions, we performed HOMER analysis on the 3xKO ATAD2 dox MC versus 3xKO dox MC. Strikingly, this revealed that the top motif enriched by ATAD2 was SOX10 itself (Fig. 4F, 4H), suggesting that ATAD2 was specifically allowing for SOX10 to bind to its target genes. Analogously, we also asked which peaks became closed after expression of ATAD2 and found that these were most highly enriched for the MITF motif (Fig. 4G, 4I), consistent with the idea that ATAD2 was associated with dedifferentiation and a decreased ability of MITF to activate target genes typically associated with differentiation. Network analysis of the loci most affected by ATAD2 and that carried the SOX10 motif showed a strong enrichment for pathways associated with neural precursor proliferation and NC migration (Fig. 4J).

### ATAD2 is necessary for NC induction

Because SOX10 is essential for proper NC induction, these data suggested that loss of ATAD2 might impair proper NC formation. To test this, we utilized the hPSC system in which we could differentiate cells into the NC in the presence or absence of sgRNAs targeting ATAD2. We utilized a previously described inducible iCas9 system (*34, 35*) to trigger the knockout during hPSC differentiation and found significant reduction with 2 different sgRNAs targeting ATAD2 confirmed by both immunofluorescence (fig. S5A and S5B) and Western blot (fig. S5C). We measured the percentage of NC cells derived from the hPSCs in this assay and found a greater than 50% reduction in NC formation with the most potent sgRNA (fig. S5D), in agreement with its requirement for activation of the SOX10 program. When taken together with the observation that the 3xKO ATAD2 dox MC become less pigmented, these data are consistent with the notion that ATAD2 facilitates access to an early NC state, in part by increasing expression of SOX10 target genes while decreasing expression of MITF target genes.

### ATAD2 forms a complex with SOX10, allowing for expression of NC genes

We next asked how ATAD2 might facilitate expression of these NC target genes. Previous work has shown that ATAD2 is able to build a protein complex together with MYC and in this way regulate a MYC-dependent signature in different cancer cell lines (*36*). We hypothesized that ATAD2 might be acting in a similar way with SOX10, by directly binding to it and facilitating transcription of its target genes. In support of this idea, we analyzed genes differentially expressed in the ATAD2^HI^ vs. ATAD2^LO^ patients from the TCGA cohort (Fig. 5A and fig. S6A). Pathway analysis revealed a strong MYC signature in the ATAD2^HI^ patients (Fig. 5A, 5C and fig. S6B), and motif analysis showed enrichment of the SOX motif (Fig. 5B). To more directly test this hypothesis, we performed co-IP experiments. As previously described, we confirmed that ATAD2 forms a complex with MYC in the 3xKO ATAD2 dox MC (Fig. 5E). Importantly, we also found that ATAD2 and SOX10 are indeed bound in the same protein complex by co-IP (Fig. 5D). Based on these findings, we hypothesized that ATAD2 might play a dual role and facilitate the expression of target genes of both MYC and SOX10 transcription factors. To test this hypothesis, we performed qRT-PCR to measure expression of representative gene targets of both of these transcription factors in the presence or absence of ATAD2 and found a significant increase in expression of key genes such as CD271, ETS1, DDX21 and E2F1. Interestingly, we also noted increases in MYC and SOX10 itself, suggesting a possible feed forward mechanism (Fig. 5F). These data indicate that ATAD2 is a critical factor that enables oncogenic gene programs by interacting with both MYC as well as the NC lineage factor SOX10.

### ATAD2 promotes melanoma phenotypes

Because SOX10 has previously been shown to be required for melanoma initiation, and its function is facilitated via ATAD2, we next wanted to test its role in melanoma initiation and progression. We first assessed in vitro cellular phenotypes using the hPSC-based system. We expressed ATAD2 in the 3xKO MC to a level similar to endogenous expression in MB cells (fig. S4A) and we analyzed their proliferation rates by incorporation of EdU followed by FACS analysis (Fig. 5G). We observed that 3xKO ATAD2 ± dox MC had a comparable proliferation rate to 3xKO dox MB and that they were significantly more proliferative than 3xKO dox MC. Both proliferation rates in 3xKO NC cells and MB increased upon BRAF^V600E^ expression, while in the already highly proliferative 3xKO ATAD2 dox MC did not. Noteworthy, in vitro 3xKO MC were not entering a senescent state upon BRAF^V600E^ expression and their proliferation rate did not change. The increased proliferation in 3xKO ATAD2 ± dox MC was accompanied by an increase in invasion, as measured by the invasion chamber analysis (fig. S7A, S7B) and as supported by ATAC-seq gene signatures consistent with an epithelial-to-mesenchymal (EMT) program (fig. S7C, S7D). We also assessed the metabolic profile of these hPSC-derived tumor lines and found evidence of significant metabolic rewiring. We used Seahorse assays to measure mitochondrial bioenergetics and glycolysis via oxygen consumption rate (OCR) and the extracellular acidification rate (ECAR). The ratio between OCR and ECAR showed that 3xKO MB were mostly relying on glycolysis for energy production and that this trend was amplified by dox-induced oncogene expression (37-39). On the contrary, 3xKO MC displayed a profoundly different metabolic profile, with sustained oxidative metabolism. Strikingly, upon ATAD2 expression, the 3xKO ATAD2 dox MC had a reduced oxygen consumption and instead switched to a more glycolytic state, as evidenced by an increased ECAR/OCR ratio (Fig. 5H and fig. S8). These phenotypes all support the notion that overexpression of ATAD2 triggers mechanisms that promote tumorigenic phenotypes.

### ATAD2 is required for melanoma initiation in vivo

Given these in vitro phenotypes and the known reliance on SOX10 for melanoma growth, these data led us to ask whether loss of ATAD2 would prevent melanoma initiation. To do this, we established a highly sensitive assay to image and quantify true tumor initiation, rather than monitoring the later steps of tumor progression. We recently developed a method called TEAZ (40) that allows us to initiate melanoma in the zebrafish skin via transgene electroporation. These transgenes can be activated in our previously developed transparent casper strain of zebrafish (41), such that we can directly visualize even a single cell as it becomes transformed by BRAF^V600E^. Into a focal region of the dorsal skin of the zebrafish, we electroporated plasmids that will activate mitfa along with BRAF^V600E^ and a GFP marker. These were co-electroporated with recombinant Cas9 as well as either non-targeting (NT) sgRNAs or sgRNAs against the zebrafish ortholog of ATAD2 (Fig. 6A). The fish were then monitored from days 3 through 21 post-electroporation, and each fish imaged using brightfield (BF) and GFP fluorescence. In the non-targeting controls, 65% of the fish developed a patch of GFP^+^ melanocytes (Fig. 6B, 6C and 6D), easily discernible from the surrounding normal skin. In contrast, in the animals which were electroporated with sgRNAs against ATAD2, we found that most fish were negative for any GFP (Fig. 6B, 6C and 6E). Quantification of the GFP^+^ area on day 14 revealed a significant decrease in overall tumor size in the ATAD2 knockout compared with the control fish (Fig. 6B). Taken together, our data supports a model in which high levels of ATAD2 expression, which is found in NC/MB cells, supports the ability of BRAF to initiate tumors by its ability to enhance lineage-specific programs in the cell of origin.

**Fig. 6.**
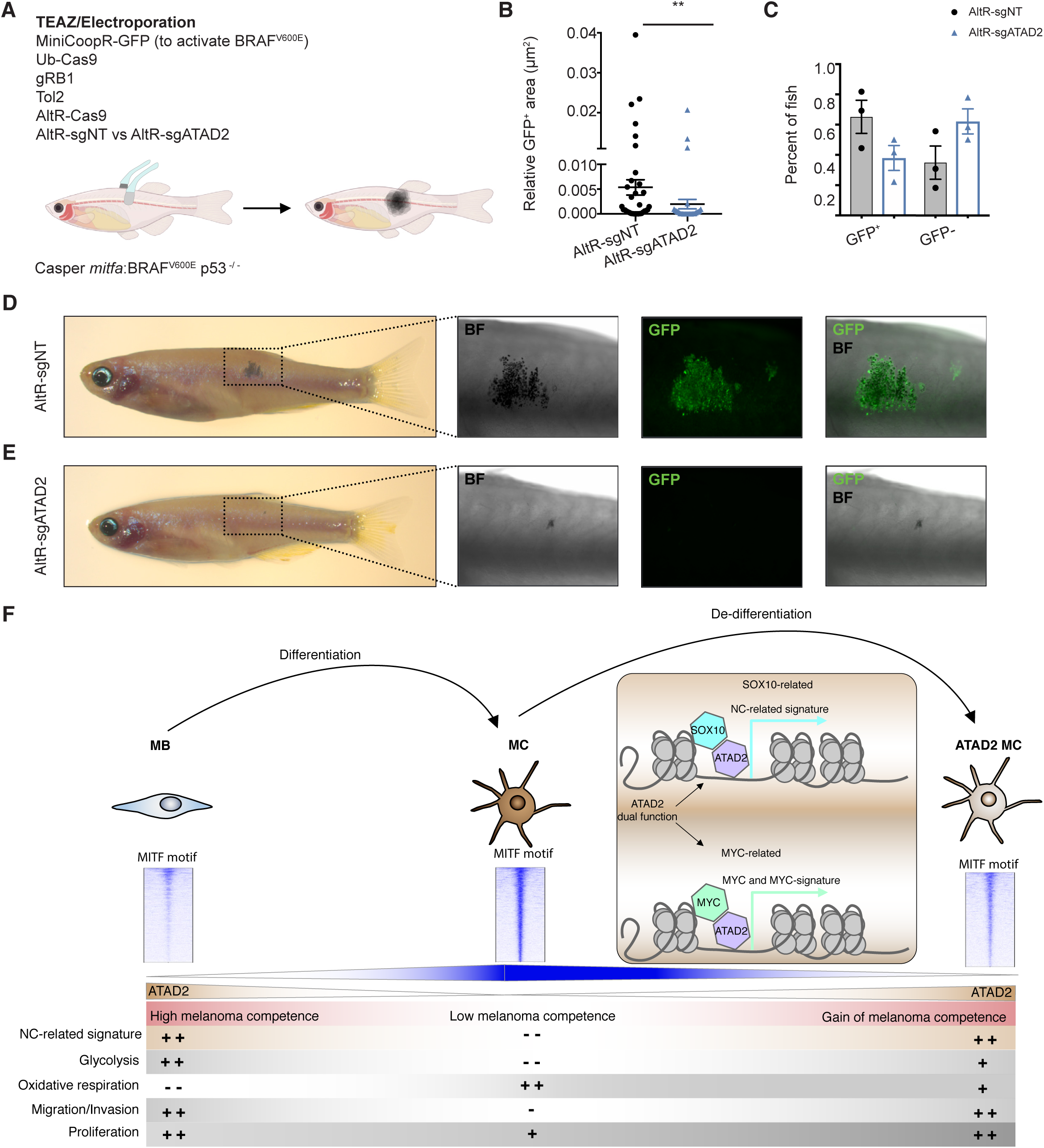
ATAD2 is required for melanoma initiation. **(A)** Schematic drawing for the TEAZ/Electroporation experiment. Fish that were p53^-/-^, mitfa^-/-^, mpv17^-/-^ and mitfa:hBRAF^V600E^ were electroporated with MiniCoopR-GFP (mitfa:MITF and mitfa:GFP), Ub-Cas9, gRB1, Tol2, AltR-Cas9, and either AltR-sgNT or a pool of AltR-sgATAD2. The fish were then monitored and quantified for melanoma initiation. **(B)** Quantification of the GFP^+^ area (µm^2^) in the transgenic fish two weeks after electroporation. Mann-Whitney test with ** p = 0.01. **(C)** Percentage of fish that were either GFP^+^ or GFP^-^ depending on the electroporation of AltR-sgNT or AltR-sgATAD2. **(D)** Representative images of a transgenic fish electroporated with AltR-sgNT. The images depict early lesions characterized by pigmentation and GFP expression. **(E)** Representative images of a transgenic fish electroporated with AltR-sgATAD2. **(F)** Summary of the roles of ATAD2 in the acquisition of melanoma competence.

## Discussion

In this study, we have established a new hPSC-derived melanoma model in concert with engineered zebrafish transgenics to investigate oncogenic competence. We showed that the ability of a cell to respond to BRAF^V600E^ depends upon the pre-existing transcriptional state of that cell, which is in turn dependent upon developmental state at the time of transformation. We find that both NC and MB stages are able to respond to BRAF^V600E^, and that ATAD2 is an oncogenic competence factor required for melanoma initiation.

NC and MB cells are most susceptible to BRAF^V600E^ and have intrinsically higher level expression of epigenetic modifiers, which include a diverse array of chromatin readers, writers and erasers. We hypothesized that this might impart these cells with a greater capability to transcriptionally respond to oncogenic insult. Consistent with this, our RNA-seq analysis demonstrated a robust response to BRAF^V600E^ only in NC and MB cells, with barely any response in the MC. This cannot be wholly explained by oncogene-induced senescence, since we find that the 3xKO MC are proliferative both before and after BRAF induction in vitro (Fig. 5G). Amongst the chromatin modifying enzymes we identified in our RNA-seq analysis, we identified ATAD2 as one example of a protein that allows for proper BRAF^V600E^-induced transformation and is amongst the most dysregulated chromatin modifying enzymes in human melanoma.

In normal physiology, ATAD2 is expressed during development in embryonic stem cells, but in adults ATAD2 expression is restricted to the male germ cells and to the bone marrow (*42*). How ATAD2 acts to affect transcription, and cancer more broadly, remains under investigation. It contains both AAA-ATPase and bromodomains and can bind to acetylated histones (*43, 44*). It has previously been shown to act as a co-regulator of oncogenic transcription factors such as MYC (*36*), and it has been identified as portending a worse prognosis in a variety of cancers (*29, 45-48*). One potential mechanism for its action may be via transcription elongation of target genes. The yeast ortholog, *Yta7*, localizes to the ORFs of highly transcribed genes (*49*) and may play a role in eviction or degradation of H3 during the elongation phase of transcription (*50*) as well as regulate the transcription of histones themselves (*51*). Our data is consistent with a model in which ATAD2 binds to the key NC factor SOX10 and allows for expression of its target genes.

One important difference between our observations and data using genetically engineered mouse models is that mature melanocytes (both zebrafish and hPSC-derived) are relatively resistant to oncogenesis. The most commonly available mouse models of melanoma utilize a *Tyrosinase*-Cre driver to activate BRAF^V600E^ (*52-55*). These animals can develop melanoma, although this can be accelerated by inactivation of tumor suppressors such as CDKN2A, TP53 or PTEN (*56-59*). Which cells within these mice act as the melanoma cell of origin has not been fully resolved (*13-15*), but our studies are not precisely comparable to the mouse studies since our zebrafish use *Tyrp1*-driven BRAF^V600E^. One possible explanation for this discrepancy is that in our system the *Tyrp1* promoter is actually driving expression in a somewhat more fully differentiated melanocyte compared to the *Tyr* promoter used in mouse studies. Another explanation is that these differences may reflect different biological thresholds for tumorigenesis, in that a different number of DNA lesions may be required to transform melanocytes in human versus mice versus zebrafish. We noted that in our hPSC-derived MC, surprisingly, even with triple knockout (of RB1, TP53, and P16), BRAF^V600E^ was still able to induce senescence in vivo. Therefore, in this particular human context these alterations are not enough to overcome oncogene-induced senescence. One possible mechanism affecting different thresholds for oncogenic competence might be the particular microenvironment, which our studies did not explicitly address. It would be important in future studies to ask how oncogenic competence might differ in each potential cell of origin depending on the local microenvironment.

Our data argue for a model in which there may not be a discrete cell of origin of melanoma, but instead support the idea that multiple cells along the spectrum from NC to MC may be capable of giving rise to tumors in the right context (Fig. 6F). It suggests that oncogenic competence is related to three interrelated factors that cooperate to determine susceptibility: DNA mutations (e.g. BRAF), cell-type specific transcription factors (e.g. SOX10), and the intrinsic levels of chromatin modifying enzymes which allow for a permissive chromatin landscape (e.g. ATAD2).

## Supporting information

Zebrafish RNA-seq

Human RNA-seq

## Acknowledgments

We are grateful to the members of the Studer laboratory and of the White laboratory for helpful discussions and support for this project. We are also thankful to the members of the Integrated Genomics Operation Core (MSKCC) for the RNA-sequencing studies, the Epigenetics Core (MSKCC) for the ATAC sequencing, the Flow Cytometry Core (MSKCC) for the cell-sorting applications, the Metabolism Core (MSKCC) for the seahorse applications, the Molecular Cytogenetics Core (MKSCC) for karyotyping and the Antitumor Assessment Core (MSKCC) for some of the xenograft studies.

This work was supported by awards from the Melanoma Research Alliance (R.M.W. and L.S.), NIH Research Program Grant R01CA229215, NIH Director’s New Innovator Award DP2CA186572, NIH Mentored Clinical Scientist Research Career Development Award K08AR055368, The Pershing Square Sohn Foundation, The Alan and Sandra Gerry Metastasis Research Initiative at the Memorial Sloan Kettering Cancer Center, The Harry J. Lloyd Foundation, Consano and the Starr Cancer Consortium (all to R.M.W.), NYSTEM award DOH01-STEM5-2016-00300 and the Starr Stem cell initiative (to L.S.) NIH Kirschstein-NRSA predoctoral fellowship F31CA196305, Joanna M. Nicolay Melanoma Foundation Research Scholar Award and the Robert B. Catell Fellowship (all to S.J.C.), Kirschstein-NRSA predoctoral fellowship F30CA220954, Melanoma Research Foundation, Medical Scientist Training Program T32GM007739 (all to N.R.C.), NIH Kirschstein-NRSA Predoctoral Fellowship F30CA236442, Molecular and Cell Biology Teaching Grant T32GM008539, Medical Scientist Training Program T32GM007739 (all to J.M.W), the Swiss National Science Foundation Early Postdoc.mobility fellowship P2ZHP3_171967 and the Postdoc.mobility fellowship P400PB_180672 (all to A.B.) and P30 CA008748 (NCI Core Facility Grant).

## Data and materials availability

GEO accession numbers for RNA-seq and ATAC-seq data are pending. All raw data files will be made available upon request. All transgenic zebrafish lines are available upon request from the authors or via the ZIRC zebrafish stock center (https://zebrafish.org/home/guide.php).

## Supplementary Materials

Materials and Methods

Figures S1-S8

References *(60-83)*

