## Supplementary Figures, Material and Methods for "Developmental chromatin programs determine oncogenic competence in melanoma"

### Materials and Methods

#### Zebrafish

*Zebrafish Husbandry*

Fish stocks were kept under standard conditions at 28.5°C under 14:10 light:dark cycles, pH (7.4), and salinity-controlled conditions. Animals were fed standard zebrafish diet consisting of brine shrimp followed by Zeigler pellets. The animal protocols described in this manuscript are approved from the Memorial Sloan Kettering Cancer Center (MSKCC) Institutional Animal Care and Use Committee (IACUC), protocol number 12-05-008.

*Transgenic lines*

Zebrafish lines used in these studies included wild-type (AB), casper (mitfa$^{-/-}$;mpv17$^{-/-}$), p53$^{-/-}$ and *mitfa*:hBRAF$^{V600E}$ line (*41*, *60-63*).

*Generating F0 transgenic lines*

One-cell-stage p53$^{-/-}$ embryos were injected with the construct Tg(promoter:BRAF$^{V600E}$-TdTomato fusion;cmlc2:eGFP) at 25 ng/µl with Tol2 mRNA at 20 ng/µl. Embryos were screened at 48 hpf for the presence of GFP in the heart as well as TdTomato in the rest of the body. Positive embryos were grown to adulthood and studied as F0 transgenics or outcrossed to identify founders that gave germline transmission of the transgene.

Cell lines

Human pluripotent stem cell (hPSC) lines employed for this study were derived from H9 (WA-09, passage 40-60) engineered to be knockout for P53, P16 and RB1 and to carry a dox-inducible expression of $BRAF^{V600E}$, passages 50-70. The pluripotent stem cells were cultured in Essential 8 (E8, Thermo Fisher Scientific) and expanded by dissociation with 0.5mM EDTA (Thermo Fisher Scientific) and plating on Vitronectin (Thermo Fisher Scientific) coated dishes. Cells were maintained at 37°C and 5% CO2 and they were routinely tested for mycoplasma and periodically assessed for genomic integrity by karyotyping. This study was approved by the Tri-institutional (MSKCC, Weill-Cornell, Rockefeller University) Embryonic Stem Cell Research Oversight (ESCRO) Committee.

Mice

NOD.Cg-$Prkdc^{scid}$ $Il2rg^{tm1Wjl}$/SzJ (NSG) mice (Jackson Laboratories) were used for the xenograft studies at the age of 3-4 months. All animal experiments were performed in accordance with protocols approved by the MSKCC IACUC. Mice were housed in rooms on a 12 h dark/light cycle at 22°C. Mice feeding was based on normal irradiated rodent diet or dox-containing diet (Envigo, 200 Dox) and water was available ad libitum. Mice were housed in groups of up to five animals per cage.

Animal experimentation

*Mice xenotransplantation*

NSG mice were injected subcutaneously with 0.5x $10^6$ hPSC-derived cells that were suspended in 50μl Growth Factor Reduced Matrigel (Corning) diluted 1:1 in DPBS (gibco). The mice were closely monitored for to primary tumor growth and for any eventual distress. 6 injections, 2 per mouse, were performed per condition.

*Zebrafish tumorigenesis assay*

Transgenic zebrafish were generated via injection into melanoma-prone ($p53^{-/-}$) one-cell embryos, and stable lines were selected by fluorescence. Fish were checked weekly for tumors formation from 4 weeks post fertilization (wpf) to 50 wpf. A Kaplan-Meier survival curve was generated using GraphPad Prism 7.

*TEAZ/Electroporation of Alt-R CRISPR/Cas9 RNP+plasmid complex into adult fish*

Alt-R Cas9 enzyme, Alt-R CRISPR-Cas9 TracrRNA, Alt-R CRISPR-Cas9 crRNAs against ATAD2 and a nontarget sequence were obtained from IDT. RNP solutions were prepared at a stock concentration of 20 µM and mixed 1:1 with the plasmid solution for a final concentration of 10 µM of RNP complex injection mix. Briefly, cRNA:TracrRNA duplexes were formed by incubating 50 µM of TracrRNA with 50 µM crRNA at 95°C for 5 minutes. TracrRNA was duplexed with either nontarget or three pooled gRNAs against ATAD2. RNP complexes were formed by mixing 50 µM Cas9 enzyme with 50 µM TracrRNA:crRNA duplex and incubated for 20 minutes at room temperature (RT). RNP duplexes were diluted to 20 µM with IDT electroporation enhancer and bought to a final concentration of 10 µM with the plasmid solution. Plasmids injected with the RNP complex included mini-CoopR-GFP (Ceol et al., 2011) (360 ng/µL), Ub-Cas9 (205 ng/µL), gRB1 (285 ng/µL) and Tol2 (80 ng/µL). Alt-R CRISPR/Cas9 RNP+plasmid complexes were electroporated into adult zebrafish (mitf-$hBRAF^{V600E}$;$mitfa^{-/-}$;$mpv17^{-/-}$;$p53^{-/-}$) as described in *40*. Adult male and female fish were anesthetized with 0.2%

tricaine and injected with 2 µL of the Alt-R CRISPR/Cas9 RNP+plasmid complex into the skin below the dorsal fin. Fish were then electroporated and returned to fresh water. Electroporation was carried out using the CM 830 Electro Square Porator from BTX Harvard Apparatus and the Genepaddles, 3×5 mm. The LV mode was used with a voltage of 45 V, 5 pulses, 60 ms pulse length and 1 s pulse interval. Electroporated zebrafish were imaged within 4 dpe to ensure successful TEAZ/electroporation, and then serially for up to 4 weeks dpe using brightfield and fluorescence imaging. Area of GFP was quantified at 2 weeks dpe using FIJI.

The gRNAs against ATAD2 were created using both GuideScan (*64*) and ChopChop (*65*) targeting the coding region of exon 1. The gRNAs used were the following:

| gRNA | Target sequence |
|---|---|
| altR_scramble | AACCTACGGGCTACGATACG |
| altR_ 001_ATAD2 | TGACAGTTTAACTAGCGCCG |
| altR_ 002_ATAD2 | CGCGGCTGAAAAAGGCCAAT |
| altR_ 003_ATAD2 | AAGCGAAAGTCCGCGCGGCT |

<u>Human PSCs culture</u>

| Small molecule | Abbreviation |
|---|---|
| SB431542 | SB |
| Y-27632 | ROCKi |
| CHIR99021 | CHIR |
| Endothelin-3 | EDN3 |
| Stem Cell Factor | SCF |

| | |
|---|---|
| Recombinant human FGF2 | FGF2 |
| Recombinant human BMP4 | BMP4 |
| Dibutyryl cAMP sodium salt | cAMP |

*Neural crest and melanoblast differentiation protocol*

Neural induction using the dual SMAD inhibition protocol was performed as previously described (*24*, *66*, *67*). Prior to start the differentiation, the hPSCs were plated as a high-density monolayer. The starting density depends on the specific hPSC line used, being WT hPSCs plated at 200,000 cells per cm$^2$ and 3xKO hPSCs plated at 100,000 cells per cm$^2$. This was required because the knockout of the tumor suppressors induced a significantly faster growth of the cells.

Day -1: Plate hPSCs on Matrigel in E8 medium with 10µM ROCKi.

Day 0-2: Change media every day with E6 media containing 1ng/ml BMP4 + 10µM SB + 600nM CHIR.

Day 2-4: Change media with E6 media containing 10µM SB + 1.5µM CHIR.

Day 4-6: Change media with E6 media containing 1.5µM CHIR.

Day 6-11: Change media every day with E6 media containing 1.5µM CHIR + 5ng/ml BMP4 100nM EDN3.

*Flow cytometry associated cell sorting*

WT and 3xKO hPSCs-derived NC and MB cells were sorted at day 11 of differentiation using a BD-FACS Aria6 cell sorter at the Flow Cytometry Core Facility of MSKCC. The cells in differentiation were initially dissociated into single cells using Accutase for 20 minutes at 37°C and then stained with conjugated antibodies against P75 (FITC anti-human CD271 (NGFR),

BioLegend) and cKIT ( Anti-Hu CD117 (cKIT) (APC), Invitrogen) to derive NC cells ($P75^+$ cells) and MB cells ($P75^+$ $cKIT^+$), and 4, 6-diamidino-2-phenylindole (DAPI) to exclude dead cells.

*Flow cytometry associated cell analysis*

iCAS9 gNT/gATAD2 ± dox hPSCs differentiated into NC cells were treated as described above and quantified for the percentage of $P75^+$ cells.

*NC cells and MB maintenance*

NC or MB spheres were maintained in ultra-low attachment plates for suspension culture.

<u>MC/MB maintenance media</u>: Neurobasal media supplemented with 1 mM L-glutamine, 0.1 mM non-essential amino acids (NEAA), N2 supplement, B27 supplement, 10ng/ml FGF2 and 3mM CHIR.

*Melanocyte differentiation*

Upon FACS sorting of $P75^+cKIT^+$ melanoblasts, 10µl cell droplets were plated onto dried PO/Lam/FN dishes. Melanocyte medium was slowly added to the plate. Continue feeding with Melanocyte medium every 2 to 3 days. Passage cells once a week at a ratio of 1:6, using Accutase for 20min at 37°C for cell detachment.

Melanocyte media: Neurobasal media + 50ng/ml SCF + 500 µM cAMP + 10ng/ml FGF2 + 3 µM CHIR + 25ng/ml BMP4 + 100nM EDN3 + B27 supplement + N2 supplement.

*hPSC AAVS1 knock-in strategy*

Knock in strategy is derived from (*34*, *68*). In 100uL of cold nucleofection solution add TALEN plasmids and Donor plasmids and mix well.

TALEN-L: 1 µg

TALEN-R: 1 µg

Puro-Donor: 5 µg

Neo-M2rtTA: 5 µg

Get single cell suspension using 0.05% Trypsin for 1 min at 37°C. Stop trypsin with DMEM/F12 10% FBS. Spin down cells at 200g X 5min. Suspend $5x10^6$ hPSCs in the nucleofector solution. Electroporate hPSCs using LONZA B16 Program in AMAXA machine. Gently plate cells at 1:2 ratio over mouse embryonic fibroblasts (MEFs) with ROCKi. 3-4 days after electroporation, add puromycin (1:2000) only for 3-4 days. One day later add geneticin (1:1000) for 4 days.

*hPSC knockout strategy*

Two guide RNAs were designed using the CRISPR design tool (http://crispor.tefor.net/) (*69*). Order gBLOCK from IDT including a U6 promoter, gRNA target and gRNA scaffold. Clone the synthesized gBlock into the empty backbone vector such as the pCR-Blunt II-TOPO. Electroporate the hPSCs with cas9-GFP fusion protein and the gRNAs. The day after, sort for $GFP^+$ cells, plate them at low density (150 cells $cm^2$) on MEFs and expand the colonies. Select for efficient knockout by sequencing and confirm it by Western Blotting.

| Name of gBLOCk | Target sequence |
|---|---|
| p53_ex5_SgRNA1 | G**CAGTCACAGCACATGACGG**AGG |

| | |
|---|---|
| p53_ex5_ASgRNA2 | A**GATGGCCATGGCGCGGACG**CGG |
| p16_ex2_ASgRNA1 | C**AGCCAGGTCCACGGGCAGA**CGG |
| p16_ex2_SgRNA2 | T**GGGCCATCGCGATGTCGCA**CGG |
| Rb1_ex1_SgRNA1 | A**GAGCAGGACAGCGGCCCGG**AGG |
| Rb1_ex1_ASgRNA3 | C**GGCGGTGCCGGGGGTTCCG**CGG |

RNA isolation and qRT-PCR

Total RNA (Trizol) was isolated using Phase Lock Gel-Heavy tubes and chloroform, followed by elution with isopropanol and washing with 75% EtOH. The RNA was purified using the RNeasy Mini Kit (Qiagen). cDNA was generated using the iScript Reverse Transcription Supermix (Bio-Rad) for RT-qPCR. For qPCR analysis, primers were obtained from QIAGEN (Quantitect Primer assays) and the reactions were performed following manufacturers' instructions using SsoFast EvaGreen®Supermix (Bio-Rad). Results were normalized to the housekeeping gene PSMB2 (*70*).

Protein isolation and Western Blotting

Cells were lysed with RIPA buffer + 1:1000 Halt™ Protease and Phosphatase Inhibitor Cocktail (Thermo Fisher Scientific) and then sonicated for 3x30sec at 4°C. Supernatant was collected upon 15min centrifugation >15000 rpm at 4°C and quantified by Precision Red Advanced Protein Assay (Cytoskeleton). Equal amounts of protein were boiled in NuPAGE LDS sample buffer (Invitrogen) at 95°C for 5 min and separated using NuPAGE 4%–12% Bis-Tris Protein Gel (Invitrogen) in NuPAGE MES SDS Running Buffer (Invitrogen). Proteins were electrophoretically transferred to a nitrocellulose membrane (Thermo Fisher Scientific) with

NuPAGE Transfer Buffer (Invitrogen). Blots were blocked for 60 min at RT in TBS-T + 5% nonfat milk (Cell Signaling) and incubated with the respective primary antibody at 4°C. The following primary antibodies were used: mouse anti-GAPDH (6C5) (Santa Cruz, 1:1000); mouse anti-Rb (IF8) (Santa Cruz, 1:1000); mouse anti-p53 (DO-1) (Santa Cruz, 1:1000); rabbit anti-CDKN2A/p16INK4A (abcam, 1:250); rabbit anti-BRAF(V600E) (sigma, 1:500); rabbit anti-p44/42 MAPK (ERK1/2) (Cell Signaling; 1:1000); rabbit anti-P-p44/42 MAPK (Cell Signaling, 1:1000); rabbit anti-ATAD2 (abcam, 1:1000). Primary antibodies were detected using the secondary anti-rabbit IgG HRP-linked (Cell Signaling, 1:1000) or the anti-mouse IgG HRP-linked (Cell Signaling, 1:1000) together with the SuperSignal™ West Femto Chemiluminescent Substrate (Thermo Fischer Scientific).

Immunostaining

Cultured cells were fixed with 4% PFA for 20min at RT, washed three times with PBS and then blocked and permeabilized in DPBS + 1% BSA and 0.2% Tween 20 for 1h at RT. Primary antibodies were diluted and incubated overnight at 4°C. The following primary antibodies were used: mouse anti-human Nanog (BD Pharmingen, 1:400); rabbit anti-ATAD2 (abcam, 1:250). Cells were subsequently washed and incubated with secondary antibodies diluted in DPBS (gibco) + 1% BSA and 0.2% Tween 20 for 1h at RT. The following secondary antibodies were used: AlexaFluor 488 Donkey anti-mouse IgG (Invitrogen, 1:500); AlexaFluor 555 Donkey anti-rabbit IgG (Invitrogen, 1:500). Finally, cells were washed 3 times with DPBS and nuclear stained with DAPI.

Histology

Zebrafish were fixed in 4% PFA for 48 hours at 4°C and then paraffin embedded. Fish were sectioned at 5 μM and placed on Apex Adhesive slides, baked at 60°C, and then stained with H&E or antibodies against GFP (Abcam, 1:100), BRAFV600E (Abcam, 1:400), phospho-RB1 (Cell Signalling, 1:400), phospho-ERK (Cell Signalling, 1:100), or SOX10 (Cell Marque, 1:50). All remaining histology of the zebrafish tumors and of the hPSC-derived tumors was performed by Histowiz (http://www.histowiz.com) and reviewed by pathologists.

Detection of proliferation

The cells were pulsed with 10 μM EdU for 30 minutes at 37°C. EdU detection was performed using the Click-iT Plus EdU Alexa Fluor 647 Flow Cytometry Assay Kit and following manufacturer's instructions (Thermo Fischer Scientific). Briefly, the cells were fixed for 15 min at RT using 100ul of Click-iT fixative per pellet and then washed with 3 mL of 1% BSA in PBS. The cells were then incubated for 30 minutes at RT in the dark. For 500μl of total reaction volume, we used 438 μl D-PBS, 10 μl Copper protectant, 2.5 μl Alexa Fluor 647 picolyl azide and 50 μl Reaction Buffer Additive. The cells were then washed with 3ml of 1x Click-iT saponin-based permeabilization and wash reagent, stained for DNA content (Hoechst) and analyzed by flow cytometer. The experiments were analyzed in the flow cytometry core facility at the MSKCC.

Oxygen Consumption Rate (OCR) and Extracellular Acidification Rate (ECAR)

We applied the XF mito stress kit using the Seahorse XFe96 Analyzer (Agilent) according to the manufacturer's protocol. In brief, we used previously coated $XF^p$ cell plates with PO/LAM/FN.

We plated out 30,000 3xKO MB cells ± dox and 10,000 3xKO MC ± dox per well and allowed them to attach for 4 hours in their respective media. The cells were then incubated with XF cell mito stress test assay medium (Agilent) + 10mM Glucose + 2 mM Glutamine + 1mM Sodium Pyruvate for 1 hour prior to the measurement in a CO2-free incubator at 37°C. During the measurement, the cells were exposed to 2.0 μM Oligomycin (Port A), 2.0 μM FCCP (Port B) and 0.5 μM Rotenone/antimycin A (Port C). Results of the measurement of both OCR and ECAR were analyzed using the Wave software (Agilent). Experiments were performed in the core facility of the Donald B. and Catherine C. Marron Cancer Metabolism Center at the MSKCC.

Invasion assay

To test the invasive properties of the hPSC-derived 3xKO ± dox MB cells, 3xKO ± dox MC and 3xKO ATAD2 ± dox MC we used the CytoSelect 24-Well Cell Invasion Assay, Basement (Cell Biolabs). We plated 0.5 x $10^6$ cells per chamber in either melanoblast media or in melanocyte media, which are both serum free. We added 500 μL of melanoblast or melanocyte media containing 10% fetal bovine serum to the lower well of the invasion plate and incubated the plate for 48h at 37°C in 5% CO2 atmosphere. Cells the crossed through the invasion chamber were stained with the Cell Stain Solution, provided with the kit (Cell Biolabs). Cell numbers were normalized taking into account the different proliferation rates using a control plate.

Cloning guides to lentiGuide-Puro vector

Cloning guides to the lentiGuide-Puro vector (Addgene, 52963) has been done following the protocol from (*35*). The guides targeting ATAD2 were designed using CRISPOR (*69*) and they were:

| gRNA | Target sequence | Oligos |
| --- | --- | --- |
| gATAD2 #1 | TAAAGCACGTGTCCGGTGGT | FWD: TAAAGCACGTGTCCGGTGGT<br>REV:<br>AAACACCACCGGACACGTGCTTTAC |
| gATAD2 #2 | ATCTTAAAGCACGTGTCCGG | FWD: ATCTTAAAGCACGTGTCCGG<br>REV:<br>AAACCCGGACACGTGCTTTAAGATC |

Viral production and viral transduction

Viruses were produced upon transfection of HEK293T cells using the Xtreme Gene 9 DNA transfection reagent (Sigma). Packaging vectors used were psPAX2 (Addgene, 12260) and pMD2.G (Addgene, 12259). The virus was collected and concentrated upon use of the centrifugal filters Amicon Ultra-15 (Millipore) 48h post-transfection. Lentiviral transduction of melanocytes was performed in medium containing viral particles overnight. 48h post transfection melanocytes were selected with antibiotics.

RNA sequencing and analysis

Total RNA was extracted as described above. Purified RNA was delivered to GENEWIZ (South Plainfield, NJ) for mRNA preparation with the TruSeq RNA V2 kit (Illumina) and 150bp paired-end sequencing on the Illumina HiSeq2500. After quality control with FASTQC (Babraham Bioinformatics) and trimming with TRIMMOMATIC (*71*), reads were aligned using STAR (*72*), with quality control via SeQC (*73*). Human PSC-derived samples were aligned to GRCh38 (Ensembl version 90) and zebrafish samples to GRCz10 (Ensembl version 81). Differential expression was calculated with DESeq2 (*74*). For zebrafish samples, pathway analysis was performed with Ingenuity Pathway Analysis (Qiagen) and with GSAA (*75*, *76*). Ortholog mapping between zebrafish and human was performed with DIOPT (*77*). In cases of more than one zebrafish ortholog of a given human gene, the zebrafish gene with the highest average expression was selected. For the hPSC-derived samples, data analysis was conducted using the Limma package, the DESeq2 package UpSETR. Human hPSC-derived samples + dox were clustered with the TCGA-SKCM primary tumor samples using the signature of melanoma subtypes published in (*28*).

ATAC sequencing (ATAC-seq)

ATAC-seq was performed as described in (*78*) with some modifications. Briefly, 50,000 freshly harvested cells were lysed in 50 µl ATAC lysis buffer (Tris 10 mM pH 7.4, 10 mM NaCl, 3 mM MgCl2, 0.1% NP-40) and incubated with Illumina Tn5 transposase at 37°C for 30 min. Samples were purified using Agencourt AMPure XP beads. Barcoding and library generation were performed using the NEBNext Q5 Hot Start HiFi PCR Master Mix with PCR amplification for

12 cycles. The size selected libraries were run on a HiSeq2500 to get an average of 40-50 million paired end reads per sample.

TCGA data analysis

Patient sample analysis was performed using data from 441 samples in the TCGA melanoma (SKCM) cancer dataset obtained from the Cancer Genome Atlas (TCGA) (*79*). We focused our analysis on only the 104 primary tumor samples (clinical data obtained from Broad Institute TCGA Genome Data Analysis Center (2019): Firehose stddata__2016_01_28 run. Broad Institute of MIT and Harvard. doi:10.7908/C11G0KM9). We used the Illumina HiSeq RNAseqv2 of the tumor samples (http://api.gdc.cancer.gov/data/3586c0da-64d0-4b74-a449-5ff4d9136611). Groups for ATAD2 expression were designated from the top and bottom quintile of RNA expression. Based on recommendations from the TCGA-CDR paper, progression free survival clinical endpoints were used for outcomes analysis (*80*). Survival analysis was performed using Kaplan Meier with log-rank p values reported. In order to be confident our clinical outcome result was not likely to be similar amongst randomly chosen genes, we randomly selected 5000 genes and repeated the same analysis. The plotted values for negative log hazards-ratios among the randomly selected genes formed a normal distribution, as expected, however ATAD2 was the highest value in the tail of the distribution, underscoring its significance in this analysis [data not shown].

It was previously shown that melanoma immune transcriptomic subtype groups are associated with different survival outcomes (Cancer Genome Atlas Network, 2015). We asked whether there was any bias in enrichment of any of the RNA subtype classes as well as the TCGA defined melanoma genomic subtypes between the defined ATAD2 expression groups and found similar distributions of the subtype classes between the ATAD2 groups suggesting clinical outcome differences were not driven by subtypes [data not shown].

For differential expression analysis (DEA), we used the 'TCGAbiolinks' R package and the TCGA Workflow to download and process Level 3 TCGA gene expression data with platform RNAseqv2 (*81*, *82*). The processing steps that were applied involve within-lane normalization procedures to adjust for GC-content effects on read counts and between-lane normalization procedures to adjust for distributional differences between lanes using the 'EDASeq' R package (*83*) as reported in (*81*, *82*). We performed a DEA between the two ATAD2 expression groups, using TCGAbiolinks, identifying differentially expressed genes (DEGs) (absolute logFC > 1, FDR <1e-05). Gene set enrichment analysis with Hallmark Pathways was performed using 'fgsea' R package and gene ontology (Biological Process) was performed using TCGABiolinks.

ATAC-seq analysis

ATAC sequencing reads were trimmed and filtered for quality and adapter content using version 0.4.5 of TrimGalore(https://www.bioinformatics.babraham.ac.uk/projects/trim_galore), with a quality setting of 15, and running version 1.15 of cutadapt and version 0.11.5 of FastQC. Reads were aligned to human assembly hg38 with version 2.3.4.1 of bowtie2 (http://bowtie-bio.sourceforge.net/bowtie2/index.shtml) and were deduplicated using MarkDuplicates in version 2.16.0 of Picard Tools. To ascertain regions of chromatin accessibility, MACS2 (https://github.com/taoliu/MACS) was used with a p-value setting of 0.001 and a publicly available melanocyte input sample (GSM3191792) was used as control. The BEDTools suite (http://bedtools.readthedocs.io) was used to create normalized read density profiles. A global peak atlas was created by first removing blacklisted regions (http://mitra.stanford.edu/kundaje/akundaje/release/blacklists/hg38-human/hg38.blacklist.bed.gz) then merging all peaks within 500 bp and counting reads with version 1.6.1 of featureCounts (http://subread.sourceforge.net). DESeq2 was used to normalize read density (median ratio method) and to calculate differential accessibility for all pairwise contrasts. Peak-gene associations were created by assigning all intragenic peaks to that gene, and otherwise using linear genomic distance to transcription start site. Gene set enrichment analysis (GSEA, http://software.broadinstitute.org/gsea) was performed with the pre-ranked option and default parameters, where each gene was assigned the single peak with the largest (in magnitude) log2 fold change associated with it. Motif signatures were obtained using Homer v4.5 (http://homer.ucsd.edu). Composite and tornado plots were created using deepTools v3.3.0 by running computeMatrix and plotHeatmap on normalized bigwigs with average signal sampled in 25 bp windows and flanking region defined by the surrounding 3 kb. Network analysis was performed using Cytoscape Enrichment Map v3.2.1 with default parameters.

Data and software availability

GEO accession numbers for RNA-seq and ATAC-seq data are pending. All raw data files will be made available upon request. All transgenic zebrafish lines are available upon request from the authors or via the ZIRC zebrafish stock center (https://zebrafish.org/home/guide.php

### Fig. S1.

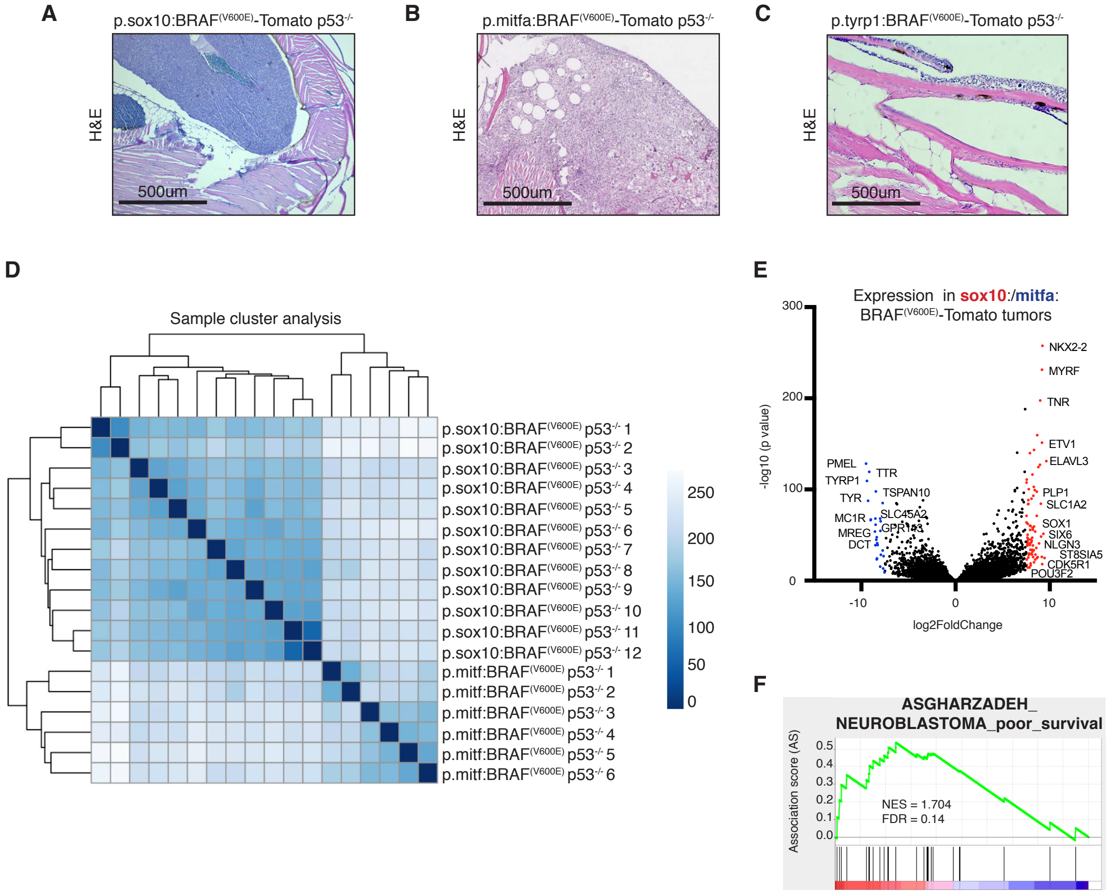


**Fig. S1. NC- and MB-derived zebrafish tumors. Related to Fig. 1.**

**(A-C)** H&E staining of NC- and MB- derived zebrafish tumors and MC-derived nevus-like structure.

**(D)** Unsupervised clustering of *sox10*:BRAF$^{(V600E)}$ p53$^{-/-}$ tumors (NC-derived; n= 12) versus *mitfa*:BRAF$^{(V600E)}$ p53$^{-/-}$ tumors (MB-derived; n= 6 biological replicates).

**(E)** Volcano plot of the human orthologs that were differentially expressed between the NC- and the MB-derived zebrafish tumors identified by RNA-seq. Top 10 most differentially expressed human orthologs in NC-derived tumors (n=6; red) and in MB-derived tumors (n=12; blue).

**(F)** GSAA showed that NC-derived zebrafish tumors expressed genes related to poor prognosis in neuroblastoma.

**Fig. S2.**

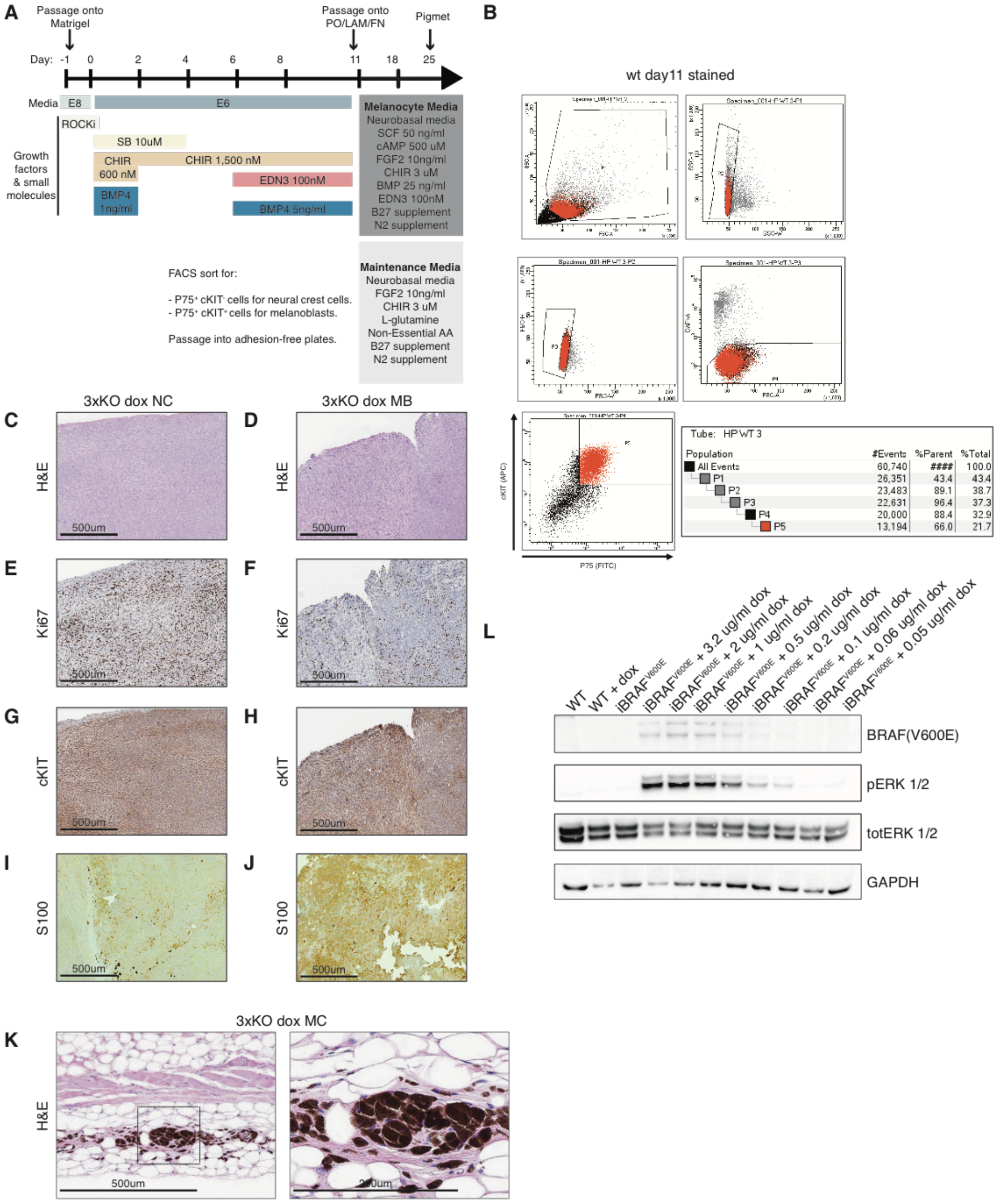

A
Passage onto Matrigel
Passage onto PO/LAM/FN
Pigmet
Day: -1 0 2 4 6 8 11 18 25
Media E8 E6
ROCKi
SB 10uM
Growth factors & small molecules
CHIR 600 nM
CHIR 1,500 nM
EDN3 100nM
BMP4 1ng/ml
BMP4 5ng/ml
Melanocyte Media
Neurobasal media
SCF 50 ng/ml
cAMP 500 uM
FGF2 10ng/ml
CHIR 3 uM
BMP 25 ng/ml
EDN3 100nM
B27 supplement
N2 supplement
FACS sort for:
- P75+ cKIT- cells for neural crest cells.
- P75+ cKIT+ cells for melanoblasts.
Passage into adhesion-free plates.
Maintenance Media
Neurobasal media
FGF2 10ng/ml
CHIR 3 uM
L-glutamine
Non-Essential AA
B27 supplement
N2 supplement
B
wt day11 stained
cKIT (APC)
P75 (FITC)
Tube: HP WT 3
Population #Events %Parent %Total
All Events 60,740 #### 100.0
P1 26,351 43.4 43.4
P2 23,483 89.1 38.7
P3 22,631 96.4 37.3
P4 20,000 88.4 32.9
P5 13,194 66.0 21.7
C
D
3xKO dox NC
3xKO dox MB
H&E
E
F
Ki67
G
H
cKIT
I
J
S100
500um
K
3xKO dox MC
H&E
500um
200um
L
WT
WT + dox
iBRAFV600E
iBRAFV600E + 3.2 ug/ml dox
iBRAFV600E + 2 ug/ml dox
iBRAFV600E + 1 ug/ml dox
iBRAFV600E + 0.5 ug/ml dox
iBRAFV600E + 0.2 ug/ml dox
iBRAFV600E + 0.1 ug/ml dox
iBRAFV600E + 0.06 ug/ml dox
iBRAFV600E + 0.05 ug/ml dox
BRAF(V600E)
pERK 1/2
totERK 1/2
GAPDH

**Fig. S2. hPSC-derived cancer model. Related to Fig. 2.**

**(A)** Schematic summary for the directed differentiation of hPSCs into NC cells, MB, and MC using a chemically defined E8/E6 media system. At day 11 of the differentiation cells were FACS sorted for P75 and cKIT expression to isolate NC cells ($P75^{+}$) and MB ($P75^{+}$ and $cKIT^{+}$). Double positive $P75^{+}$ $cKIT^{+}$ MB were further differentiated into mature MC.

**(B)** FACS sorting for P75 (FITC) and cKIT (APC) double positive cells at day 11 of the differentiation.

**(C-J)** Immunohistochemistry staining for H&E, Ki67, cKIT and S100 in 3xKO dox NC-derived and 3xKO dox MB-derived tumors.

**(K)** Immunohistochemistry staining for H&E in 3xKO dox MC that built a nevus-like structure upon transplantation.

**(L)** Western blot analysis for the dox-inducible expression of $BRAF^{(V600E)}$ and the following activation of the MAPK pathway upon 72h dox treatment.

**Fig. S3.**

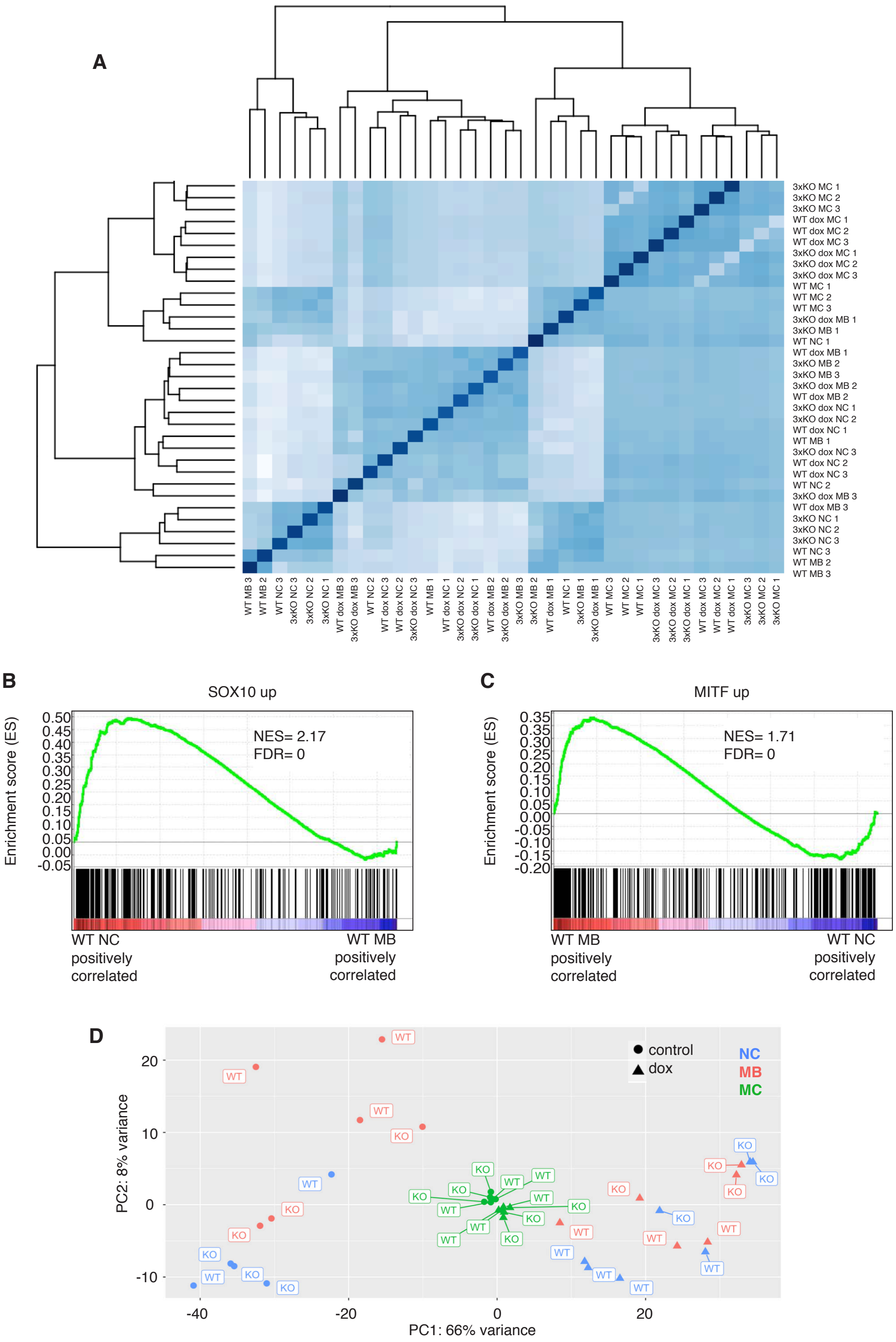
A
3xKO MC 1
3xKO MC 2
3xKO MC 3
WT dox MC 1
WT dox MC 2
WT dox MC 3
3xKO dox MC 1
3xKO dox MC 2
3xKO dox MC 3
WT MC 1
WT MC 2
WT MC 3
3xKO dox MB 1
3xKO MB 1
WT NC 1
WT dox MB 1
3xKO MB 2
3xKO MB 3
3xKO dox MB 2
WT dox MB 2
3xKO dox NC 1
3xKO dox NC 2
WT dox NC 1
WT MB 1
3xKO dox NC 3
WT dox NC 2
WT dox NC 3
WT NC 2
3xKO dox MB 3
WT dox MB 3
3xKO NC 1
3xKO NC 2
3xKO NC 3
WT NC 3
WT MB 2
WT MB 3
B
SOX10 up
Enrichment score (ES)
NES= 2.17
FDR= 0
WT NC positively correlated
WT MB positively correlated
C
MITF up
Enrichment score (ES)
NES= 1.71
FDR= 0
WT MB positively correlated
WT NC positively correlated
D
control
dox
NC
MB
MC
PC2: 8% variance
PC1: 66% variance

**Fig. S3. Cancer competent NC/MB states show a transcriptional response to oncogene activation. Human PSCs-derived NC cells and MB resemble the NC/MB-derived tumors in the zebrafish. Related to Fig. 2.**

**(A)** Unsupervised clustering of WT ± dox NC, MB and MC and 3xKO ± dox NC, MB and MC.

**(B)** GSEA of NC-derived zebrafish tumors versus WT hPSC-derived NC cells showed a similar transcriptional profile.

**(C)** Similarly, GSEA of MB-derived zebrafish tumors versus WT hPSC−derived MB showed a similar transcriptional profile.

**(D)** PCA of the RNA-seq data generated from all the samples shows transcriptional response of the NC/MB states to oncogene activation.

**Fig. S4.**

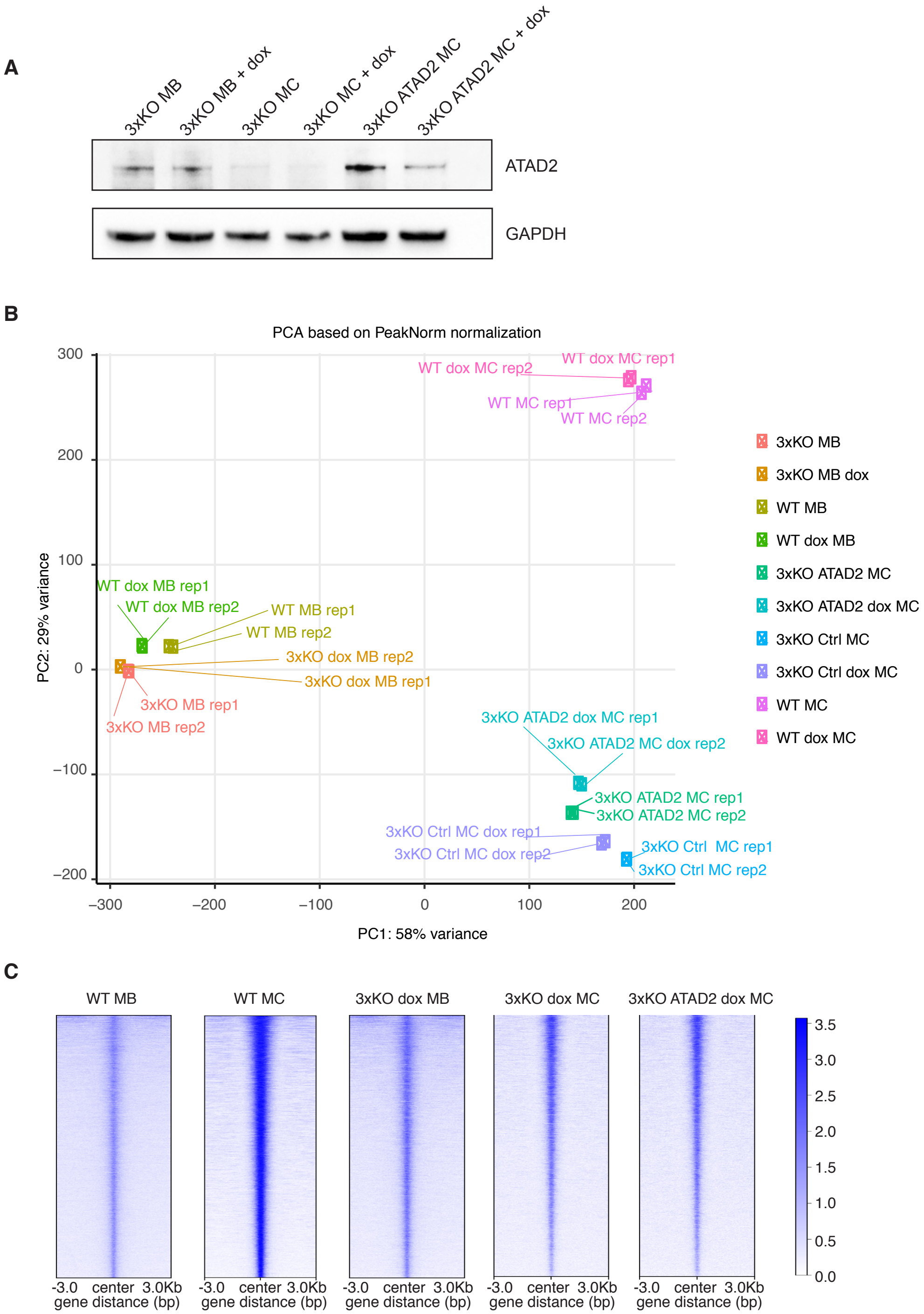
A
3xKO MB
3xKO MB + dox
3xKO MC
3xKO MC + dox
3xKO ATAD2 MC
3xKO ATAD2 MC + dox
ATAD2
GAPDH
B
PCA based on PeakNorm normalization
PC2: 29% variance
PC1: 58% variance
300
200
100
0
−100
−200
−300
−200
−100
0
100
200
WT dox MC rep1
WT dox MC rep2
WT MC rep1
WT MC rep2
WT dox MB rep1
WT dox MB rep2
WT MB rep1
WT MB rep2
3xKO dox MB rep2
3xKO dox MB rep1
3xKO MB rep1
3xKO MB rep2
3xKO ATAD2 dox MC rep1
3xKO ATAD2 MC dox rep2
3xKO ATAD2 MC rep1
3xKO ATAD2 MC rep2
3xKO Ctrl MC dox rep1
3xKO Ctrl MC dox rep2
3xKO Ctrl MC rep1
3xKO Ctrl MC rep2
3xKO MB
3xKO MB dox
WT MB
WT dox MB
3xKO ATAD2 MC
3xKO ATAD2 dox MC
3xKO Ctrl MC
3xKO Ctrl dox MC
WT MC
WT dox MC
C
WT MB
WT MC
3xKO dox MB
3xKO dox MC
3xKO ATAD2 dox MC
-3.0 center 3.0Kb
gene distance (bp)
3.5
3.0
2.5
2.0
1.5
1.0
0.5
0.0

**Fig. S4. ATAD2 expression reactivates specific programs related to development, without causing an overall reprogramming. Related to Fig. 4.**

**(A)** Western blot analysis of ATAD2 protein levels in 3xKO MB ± dox, 3xKO MC ± dox and 3xKO ATAD2 MC ± dox, 72h dox treatment.

**(B)** PCA of the ATAC-seq of WT ± dox MB and WT ± dox MC, 3xKO ± dox MB and 3xKO ± dox MC and 3xKO ATAD2 ± dox MC.

**(C)** Tornado plots of the ATAC-seq of WT MB, WT MC, 3xKO dox MB, 3xKO dox MC and 3xKO dox ATAD2 MC.

Fig. S5.

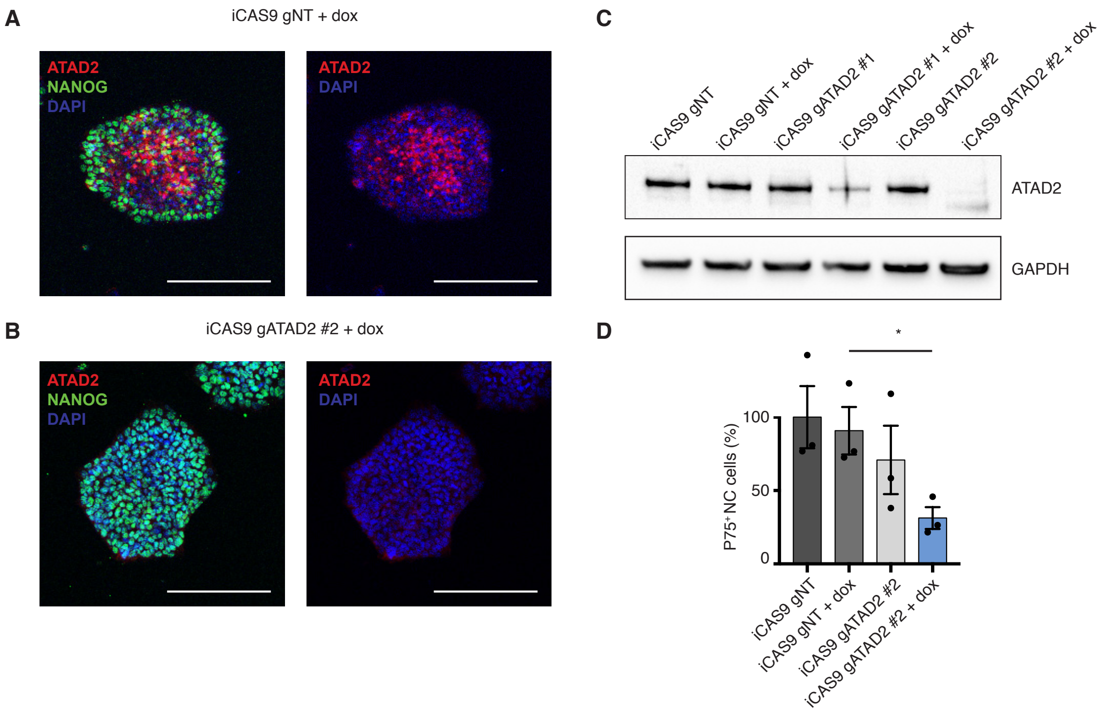


**Fig. S5. ATAD2 is required for proper NC formation. Related to Fig. 4.**

**(A-B)** Immunostaining for ATAD2 and NANOG in iCAS9 gNT hPSCs **(A)** and in iCAS9 gATAD2 hPSCs **(B)** + 72h dox treatment.

**(C)** Western blot for ATAD2 and GAPDH on protein lysates of iCAS9 gNT or gATAD2 hPSCs ± 72h dox treatment.

**(D)** FACS quantification of total number of NC cells (p75$^{+}$ cells) during the differentiation of iCAS9 gNT/gATAD2 hPSCs ± dox treatment. Unpaired t test with * = $p < 0.05$.

**Fig. S6.**

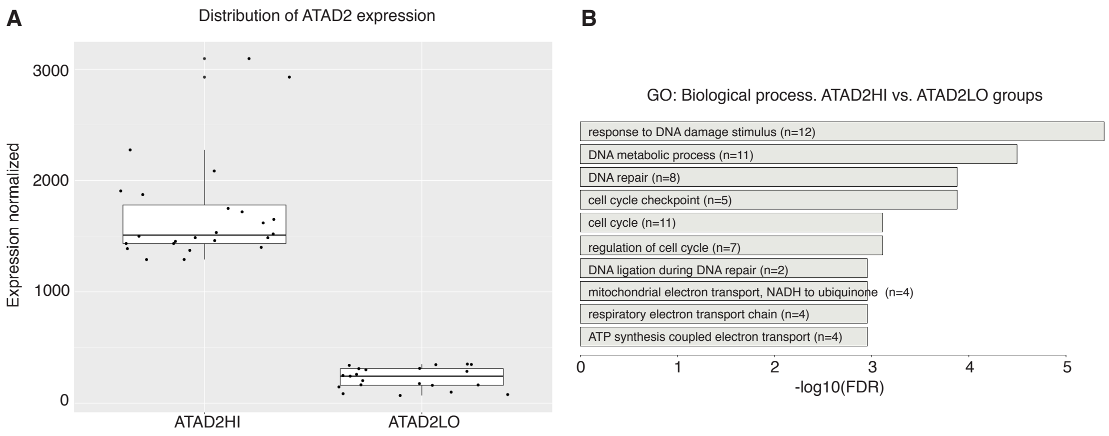


**Fig. S6. Clinical data related to ATAD2 expression in TCGA SKCM patients. Related to Fig. 3 and Fig. 5.**

**(A)** ATAD2 mRNA expression profile in primary cutaneous melanomas and separation between ATAD2 high (ATAD2HI) mRNA level patient group and ATAD2 low (ATAD2LO) mRNA level patient group.

**(B)** GO analysis of biological processes in the ATAD2HI patient group compared to the ATAD2LO patient group.

**Fig. S7.**

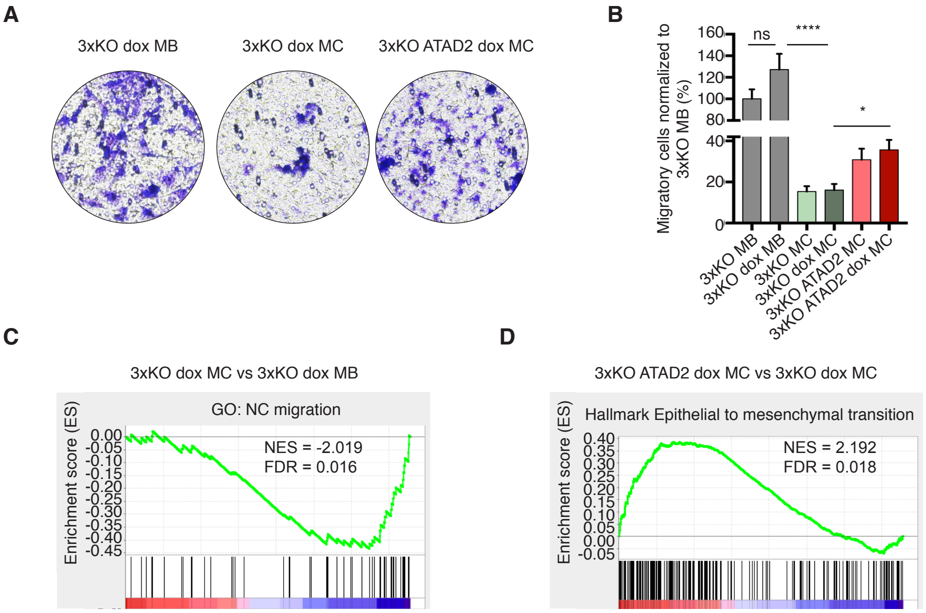


**Fig. S7. MB cells are invasive cells and ATAD2 expression in 3xKO MC induces an invasive phenotype. Related to Fig. 5.**

**(A, B)** Representative pictures **(A)** and quantifications **(B)** of the 48h invasion assay for 3xKO ± dox MB, 3xKO ± dox MC and 3xKO ATAD2 ± dox MC. Data are shown as mean ± SEM normalized to cell numbers after 48h, n=3. * = $p < 0.05$; ** = $p < 0.0001$.

**(C)** GSEA of the ATAC–seq comparing 3xKO dox MB over 3xKO dox MC shows an enrichment for a signature related to NC migration.

**(D)** GSEA of the ATAC–seq comparing 3xKO ATAD2 dox MC over 3xKO dox MC shows an enrichment for a signature related to an epithelial to mesenchymal transition.

**Fig. S8.**

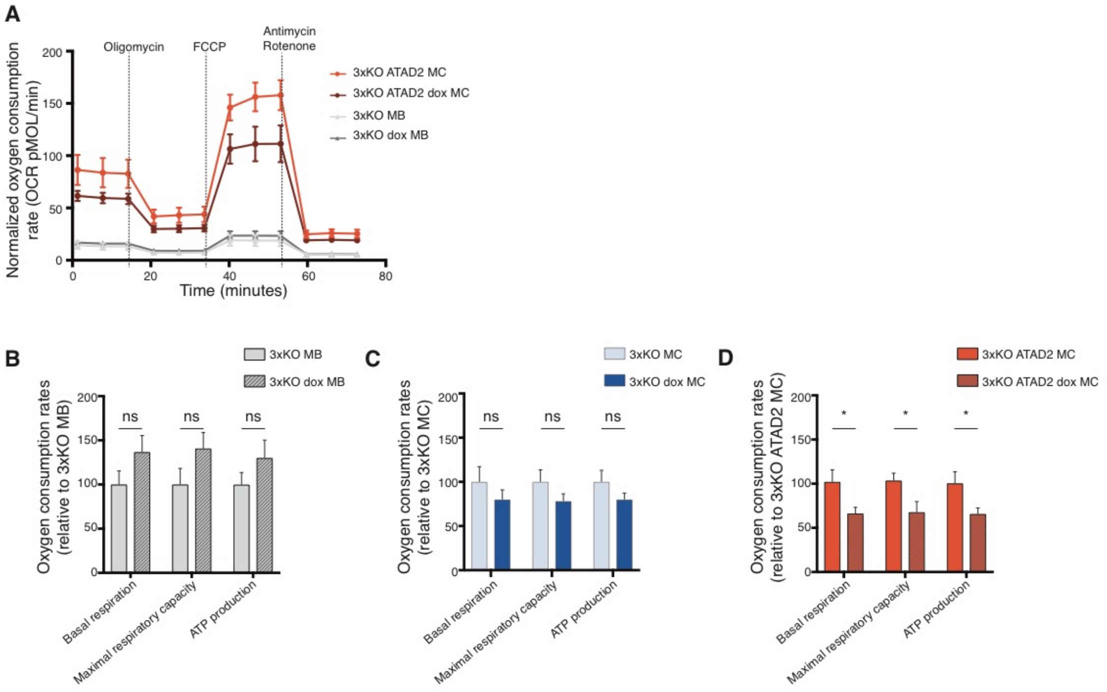


**Fig. S8. ATAD2 regulates a metabolic response upon BRAF$^{V600E}$ expression. Related to Fig. 5.**

**(A-D)** OCR measurements **(A)** and mitochondrial bioenergetics **(B-D)** in 3xKO MB ± dox **(B)**, in 3xKO MC ± dox **(C)** and 3xKO ATAD2 MC ± dox **(D)**, with 72h dox treatment. Data are shown as mean ± SEM; n = 3. p values were calculated by unpaired Student's t test. * = $p < 0.05$.
